# PhylEx: Accurate reconstruction of clonal structure via integrated analysis of bulk DNA-seq and single cell RNA-seq data

**DOI:** 10.1101/2021.02.16.431009

**Authors:** Seong-Hwan Jun, Hosein Toosi, Jeff Mold, Camilla Engblom, Xinsong Chen, Ciara O’Flanagan, Michael Hagemann-Jensen, Rickard Sandberg, Samuel Aparicio, Johan Hartman, Andrew Roth, Jens Lagergren

## Abstract

We propose PhylEx: a clonal-tree reconstruction method that integrates bulk genomics and single-cell transcriptomics data. In addition to the clonal-tree, PhylEx also assigns single-cells to clones, which effectively produce clonal expression profiles, and generates clonal genotypes. By analyzing scRNA-seq integrated with bulk DNA-seq, PhylEx can take advantage of co-occurrences of the mutations found in the cells. In the probabilistic model underlying PhylEx, the raw read counts from scRNA-seq follow a mixture of Beta-Binomial distributions, which accounts for the sparse nature of single-cell gene expression data; the mixture lessens the penalty caused by mutations not observed due to mono-allelic expression. We rigorously evaluated PhylEx on simulated datasets as well as a biological dataset consisting of a previously well-characterized high-grade serous ovarian cancer (HGSOC) cell line. PhylEx outperformed the state-of-the-art methods by a wide margin both when comparing capacity for clonal-tree reconstruction and capacity for correctly clustering mutations. By analyzing HGSOC and HER2+ breast cancer data, we also show that PhylEx clears the way for phylo-phenotypic analysis of cancer, i.e., that the clonal expression profiles, induced by the cell-to-clone assignments, can be exploited in a manner beyond what is possible with only expression-based clustering.

## Introduction

Cancer is an evolutionary process with the ongoing mutational processes coupled with selection and drift leading to genetic diversity within the tumour cell populations. Though each cell is fundamentally distinct in a cancer, there typically exist groups of cells which are genomically nearly identical, so called clonal populations [1]. The evolutionary relationship between clones can be represented by a phylogenetic tree or clonal-tree. Inferring clonal population structure, genotypes and trees from sequence data has become an active area of research in the past decade. Early approaches used bulk sequence data coupled with computational deconvolution to address the admixed nature of bulk data [2, 3, 4, 5, 6]. Recent advances in single cell DNA (scDNA) sequencing technologies have prompted the development of approaches better tailored to these data types [7, 8, 9, 10, 11].

Though the aforementioned methods can resolve clonal population structure, they cannot identify functional differences that result from the genomic heterogeneity. The increasing availability of single cell RNA (scRNA) sequencing data provides a promising approach to address this problem. Recent methods that seek to assign gene expression profiles to clones have treated the problem as a two step procedure whereby the clonal population structure is identified and then scRNA data is aligned to clonal genotypes [12, 13]. This two stage approach is potentially statistically inefficient as information in the scRNA data cannot be used to improve clonal population structure. Hence, there is an unmet need for integrative approaches to jointly analyse DNA and scRNA data to simultaneously identify clonal population structure and the associated clonal gene expression profiles.

In this work we consider the problem of performing simultaneous inference of clonal-trees, genotypes, and expression profiles by jointly analyzing bulk DNA and scRNA sequencing data. We introduce a Bayesian probabilistic method called PhylEx to solve this problem. PhylEx leverages information about the SNVs observed within a single cell to both improve clonal-tree reconstruction, and to assign RNA expression profiles to clones.

The method takes as input allelic count data from both DNA and scRNA but also copy number data from DNA. The outputs of the method are: a clonal-tree, a genotype defined by the presence or absence of SNVs for each clone, and an assignment of each cell with scRNA data to a clone. Using this information, we can aggregate the scRNA data by clone to infer clonal, pseudo bulk based, gene expression profiles for further analysis.

There have been several approaches which have considered integrating single cell and bulk sequencing data. The most closely related to PhylEx is ddClone which uses both bulk and scDNA data [10]. In contrast to our approach ddClone does not infer a clone tree. Additionally ddClone cannot infer clonal gene expression profiles because RNA data is not used. Also closely related to PhylEx are approaches which consider the problem of assigning scRNA data to clones using known clonal genotypes. The earliest approach we are aware of is clonealign which uses clonal copy number profiles and scRNA data to generate clonal expression profiles [12]. In the original publication the copy number profiles were inferred from scDNA data, though in principle they could also be inferred from bulk sequencing. In contrast to PhylEx, clonealign does not infer a phylogeny. Furthermore, clonealign requires that there is sufficient copy number variability between clones to uniquely correlate scRNA expression to genotypes. Thus clonealign is not applicable to cancers without significant copy number variation. Cardelino is another approach for aligning scRNA and clonal genotypes which uses SNV data [13]. In contrast to PhylEx, Cardelino assumes that (1) the clonal-tree is known and remains fixed during inference and (2) that the genotypes are known and remains more or less fixed during inference. As a result Cardelino is sensitive to the clonal information input, which can be noisy if inferred purely from bulk DNA.

In the remainder of this work we describe the PhylEx probabilistic model and the computational techniques used for parameter inference. Next we systematically benchmark our approach using synthetic data and compare it to existing state-of-the-art clone reconstruction methods. We then evaluate the performance of PhylEx by analyzing Smart-Seq3 from cell lines thoroughly investigated using the DLP approach, in [14]. Finally, we apply PhylEx to novel breast cancer Smart-Seq3 data to study the clonal expression profiles.

## Results

### Method overview

PhylEx is a Bayesian statistical tool that simultaneously reconstructs a clonal-tree and assigns single-cells, as well as genotypes, to the clones for a tumor characterized by bulk DNA-seq and scRNA-seq data (Figure 1 **a**). The standard bulk data processing is performed, including variant and copy number calls to identify loci with SNVs and their copy number profiles (Supplement). For each locus, the bulk data consists of the number of reads mapping to the variant allele and the total number of reads mapping to the locus. Similarly, standard scRNA-seq data processing is applied to align and map the reads for each cell, yielding data that consists of the total depth and the number of reads mapping to the variant allele for each locus. The underlying statistical model is based on the tree-structured stick breaking process, a flexible prior distribution over the clonal-tree structure [15], and an infinite site model that define a distribution over clonal genotypes. The model has an observational component for the single-cell expression and the DNA-seq data given the clonal-tree and the genotypes associated with the clones. The inference machinery takes advantage of slice sampling to explore the space of clonal-trees [16] and Metropolis-Hastings for exploring the clone fractions [3, 4].

**Figure 1:**
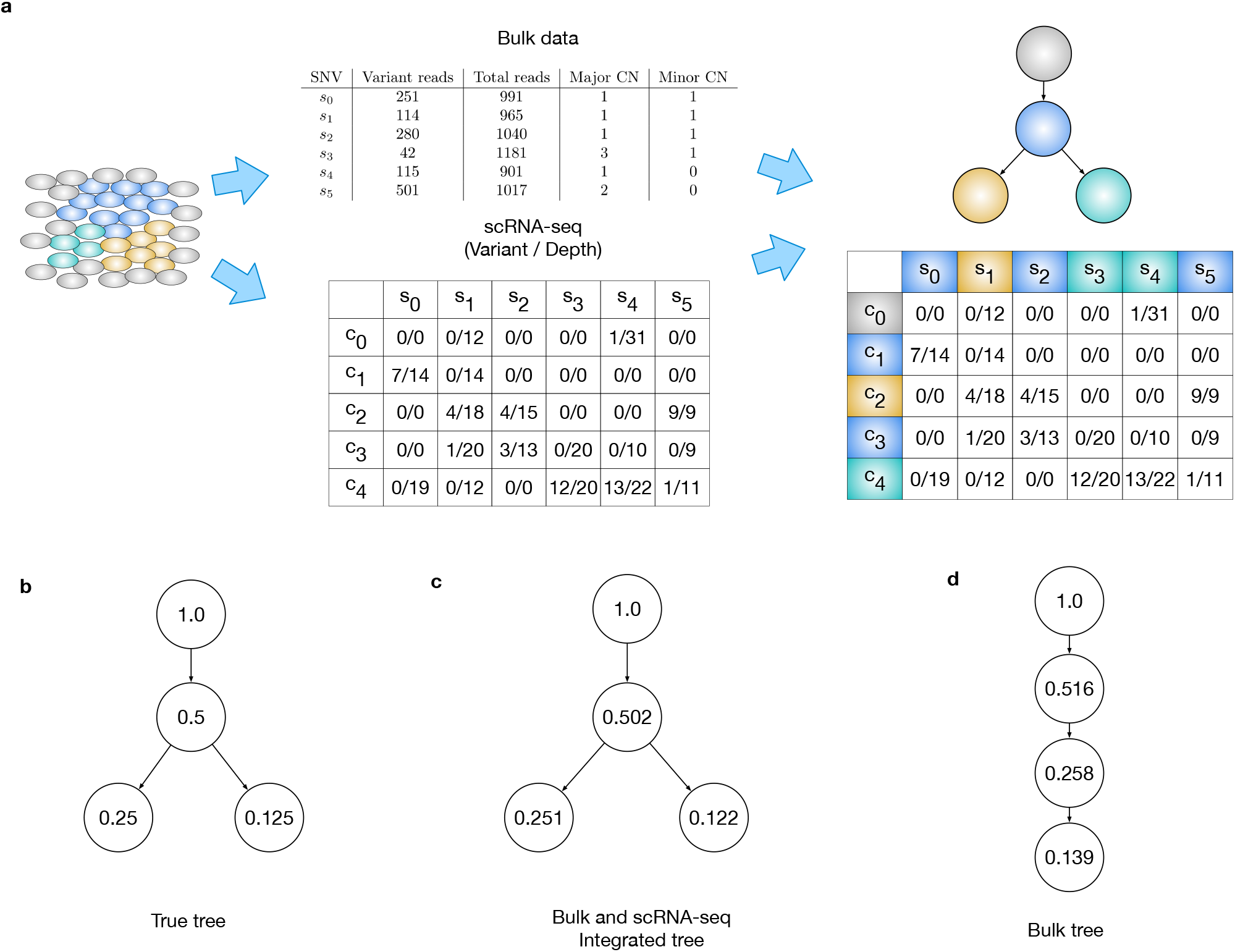
a. Schematic diagram describing the bulk DNA-seq and scRNA-seq data input. The output of PhylEx includes the tree and assignment of SNVs and cells to clones. b. The cherry shaped tree used in the illustrative example for identifying branching structure from scRNA-seq. The cellular prevalences are indicated for each clone. c. Inferred tree from integrated analysis of bulk DNA-seq and scRNA-seq. d. Tree inferred using bulk DNA-seq.

The observational model specifies that the read counts from the bulk data follow a binomial distribution. The probability of observing a variant read is a function of the unobserved cellular prevalence and an estimated clonal copy-number (Method), and its definition follows that made for clonal analysis based solely on bulk DNA-seq data [2, 3]. The observational model for the scRNA-seq data is a mixture of two Beta-Binomial distributions: one for the monoallelic and one for the biallelic expression. The scRNA-seq gene expression data frequently exhibit bursty expression [17]. The mixture distribution functions to lessen the penalty of assigning a cell which does not express a gene with a mutation to a clone with a genotype that has the mutation. PhylEx marginalizes over all possible cell-to-clone assignments to evaluate the likelihood of the single-cell data. The marginalization has the positive effect of removing uncertainty in scoring the clonal-tree due to uncertain cell-to-clone membership.

PhylEx generates samples from the posterior distribution over the clonal-tree as well as a *maximum a posteriori* (MAP) tree. The output also includes clonal genotypes and cell-to-clone assignments. The clone analysis conducted by PhylEx then facilitates a range of differential expression investigations on the otherwise inaccessible tumor clones.

### Integrating scRNA with bulk DNA improves clonal-tree reconstruction

We begin with an illustrative example to test the strength of the co-occurrence signal in single-cell data. We simulated bulk and scRNA-seq data for 100 SNVs and 20 single-cells over a cherry shaped tree (Figure 1 **b**) under an evolutionary model devoid of copy-number aberrations. We analyzed this data using PhylEx and PhyloWGS [4]. Figure 1 **c, d** show the maximum *a posteriori* (MAP) trees from PhylEx and PhyloWGS respectively. Both methods infer the cellular prevalences correctly, but the tree inferred by PhyloWGS incorrectly has a linear topology. The linear and cherry tree explain the observed variant allele frequencies (VAFs) equally well in the sense they have equal data likelihoods. This example highlights that estimating clonal-trees from single sample bulk DNA data is an unidentifiable problem. PhylEx correctly infers the clonal-tree by taking advantage of the co-occurrence of mutations in the single-cell data and performs co-clustering of the SNVs and cells (Supplementary Figure 1).

We performed a comprehensive study of simulated data on larger trees and a model of evolution involving copy-number changes. Copy-number variation obfuscate the VAF, which renders bulk data-based clonal-tree reconstruction an underdetermined problem. We used two clonal-tree reconstruction methods, PhyloWGS and Canopy, for comparison. PhyloWGS requires subclonal copy number calls as an input; since such data is not available for simulated data, we implemented the methodology underlying PhyloWGS, which we refer to as TSSB, to investigate the performance of the PhyloWGS methodology. Canopy [6] is a Bayesian clonal-tree reconstruction software that takes advantage of clonal copy-number information.

As the cancer evolution can involve multifurcating events [18], we simulated the data using multifurcating trees and a binary tree with a fixed depth (Supplement). We found that for both binary and multifurcating trees, PhylEx outperformed Canopy and PhyloWGS/TSSB in all clustering metrics (Supplementary Figures 2 and 3), and the ancestral reconstruction error (Figure 2 **a, b**). The performance of PhylEx improves progressively with the number of cells, as hoped. Comparing PhylEx to bulk-based clonal-tree reconstruction methods further demonstrates that scRNA-seq data can mitigate the negative impact of copy-number changes on clonal-tree reconstruction accuracy.

**Figure 2:**
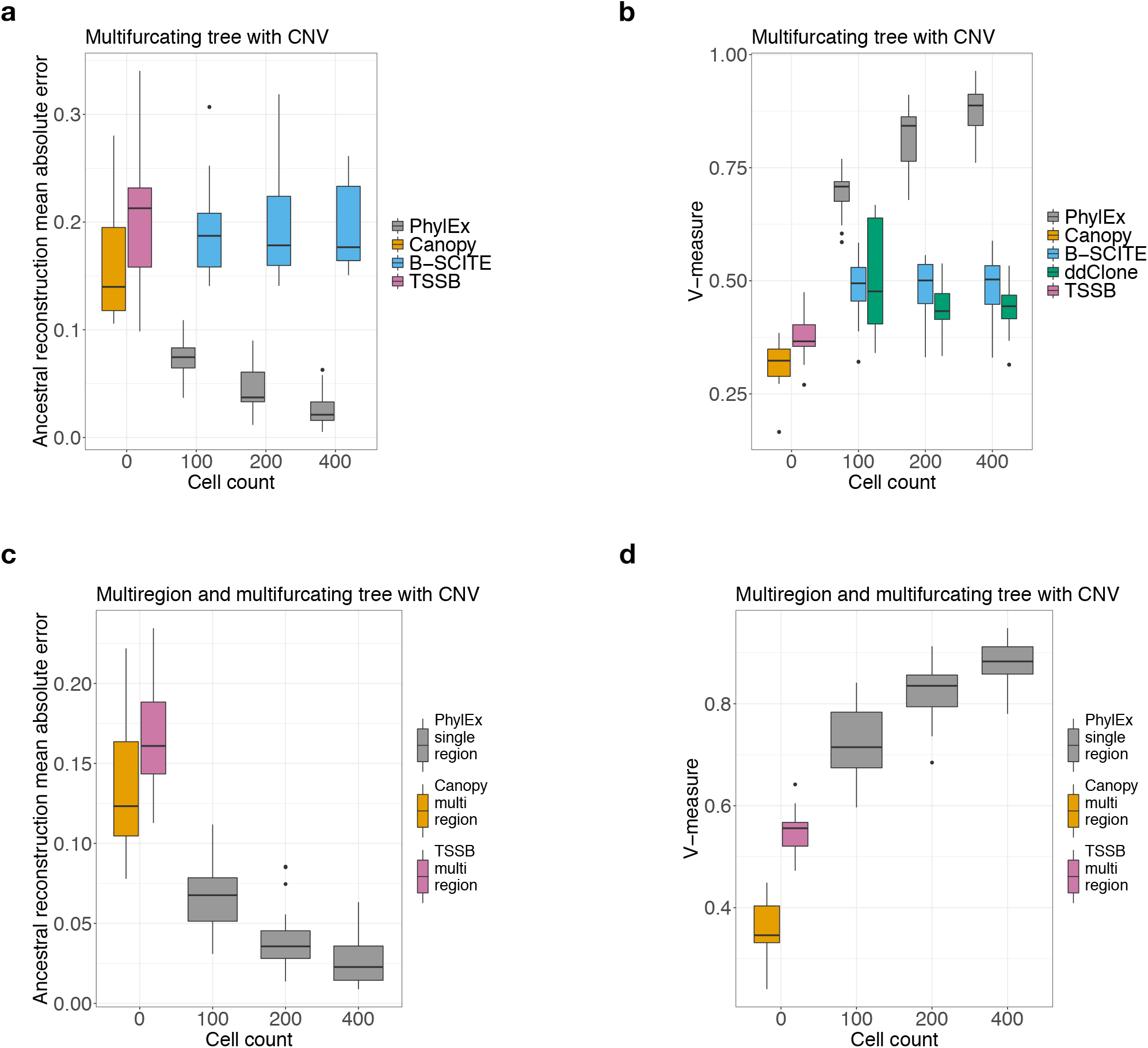
Simulated data analysis results with *n* = 20 data replicates generated with 100 SNVs in each replicate from multifurcating tree with copy number evolution scenario. a. Compared PhylEx to Canopy, TSSB, and B-SCITE on tree reconstruction error and b. on V-measure clustering metric. c-d. Comparison of PhylEx using single-region bulk DNA-seq and scRNA-seq to bulk based methods supplied with multi-region DNA-seq; tree reconstruction error and V-measure clustering metric are used in the comparison.

### Single-region bulk-seq with scRNA-seq outperforms multi-region bulk-seq

Multi-region sequencing is a standard approach to improve the accuracy of the clonal-tree reconstruction, e.g., to resolve branching better [2, 3, 4, 19]. For solid tumors, spatial samples are taken as statistical replicates with common evolutionary history but possibly with different cellular prevalences. However, depending on the type of tumor, spatial sampling may not be feasible. In particular, multi-regional sampling is difficult to perform without prior surgical tumor removal, preventing it from being viable for pre-surgical treatment decisions. We evaluated the performance of PhylEx on data consisting of a *single-region* bulk DNA-seq combined with scRNA-seq data against bulk methods supplied with multi-region DNA data.

We used a multifurcating tree and simulated the bulk DNA data both under one model with and one model without copy-number evolution. Devoid of copy number evolution and given multi-region data, the bulk methods achieved high accuracy (Supplementary Figure 4): for example, PhyloWGS and TSSB achieved 0.85 in the V-measure metric on multifurcating trees (Supplementary Figure 5). Nevertheless, when supplied with single-cell data, PhylEx performed better, achieving a V-measure metric upwards of 0.95 using 400 cells on all evaluation metrics (Supplementary Figure 4). With data simulated under a copy-number evolution model, bulk clonal-tree reconstruction methods struggled even when supplied with multi-region data. On the contrary, PhylEx improved the accuracy given only a single-region bulk DNA-seq data by integrating scRNA-seq data in the analysis (Figure 2 **c, d**, Supplementary Figure 6). This investigation demonstrates that researchers can reconstruct high-quality clonal-trees using single region bulk DNA and scRNA sequencing.

### Specialized method to integrate bulk and scRNA-seq are necessary

We compared PhylEx to two methods for integrating bulk DNA-seq and scDNA-seq, B-SCITE and ddClone [10, 11]. One of the challenges of using these scDNA-seq methods is that they require a variant calling as a pre-processing step, i.e., for each cell and each locus, determine the presence or absence of a mutation. Although variant calling is an active field of research, it remains a challenging problem with the potential for high false positive (FP) and false-negative (FN) rates. Adopting these methods for scRNA-seq is sure to suffer from FP and FN problems as the expression profile is inherently sparse, bursty, with frequent monoallelic expression. A key feature of PhylEx is that it works directly with the read counts and does not require variant calling.

We found that PhylEx outperformed both of these methods on synthetic data generated from both binary and multifurcating trees, under evolutionary models with and without copy numbers aberrations (Figure 2 **a, b**, Supplementary Figures 2, 3, 4). Furthermore, PhylEx exhibited an increase in performance with an increasing number of cells. In contrast, the other methods did not benefit from having more cells, likely because having more cells implies a higher incidence of FP and FN variant calls. Our results suggest that specialized methods for integrating bulk genomics with single-cell transcriptomics are needed to extract the signal from scRNA-seq data.

### PhylEx resolves high-grade serous ovarian cancer cell-line

To assess the performance of PhylEx on real data, we analyzed a set of high-grade serous ovarian cancer cell-line which have recently had their clonal structure accurately determined using scDNA sequencing [14]. These cell-lines are derviced from the same patient, one from the primary tumor (OV2295) and two from relapse specimens (OV2295R2 and TOV2295R). The clonal-tree for these cell-lines was inferred using a combination of copy-number, structural variants and SNVs identified using scDNA data. Consequently, by considering their clonal-tree and their assignment of SNVs to the clones as *de facto ground truth*, we could evaluate the performance of PhylEx on a realistic biological data set.

We performed Smart-Seq3 scRNA-seq [20] on OV2295 and OV2295R2^1^. We constructed a single-region pseudo-bulk data by combining the scDNA from the two regions. We obtained 360 scRNA-seq cells passing quality control and identified 67 SNVs with coverage in the scRNA data. We excluded 19 SNVs from performance evaluation due to the incompatible annotation of SNV-to-clone assignment published in [14] resulting in 47 SNVs used for testing. However, we used all 67 SNVs when fitting PhylEx.

There is strong concordance between the *ground truth* clonal-tree and the PhylEx MAP clonal-tree. First, when disregarding a node, labelled *EFGHI*, of the *ground truth* clonal-tree with a single SNV, these trees have the same topology (Figure 3 **a, b**, Supplementary Table 1). PhylEx correctly assigned 23 of 24 ancestral mutations. One SNV in the *ground truth ABCD* clone was mistakenly assigned to the *ground truth CD* clone. The clones *EF* and *EFGHI* were clumped together, thereby also incorrectly clustering the single SNV in *EFGHI* along with the SNVs in clone *EF*. We compared the results of PhylEx to those inferred with Canopy, TSSB, ddClone, and B-SCITE [6, 3, 4, 10, 11]. To quantify performance, we used the same metrics as in the synthetic data experiments. PhylEx outperformed all of the other methods (Table 1).

**Figure 3:**
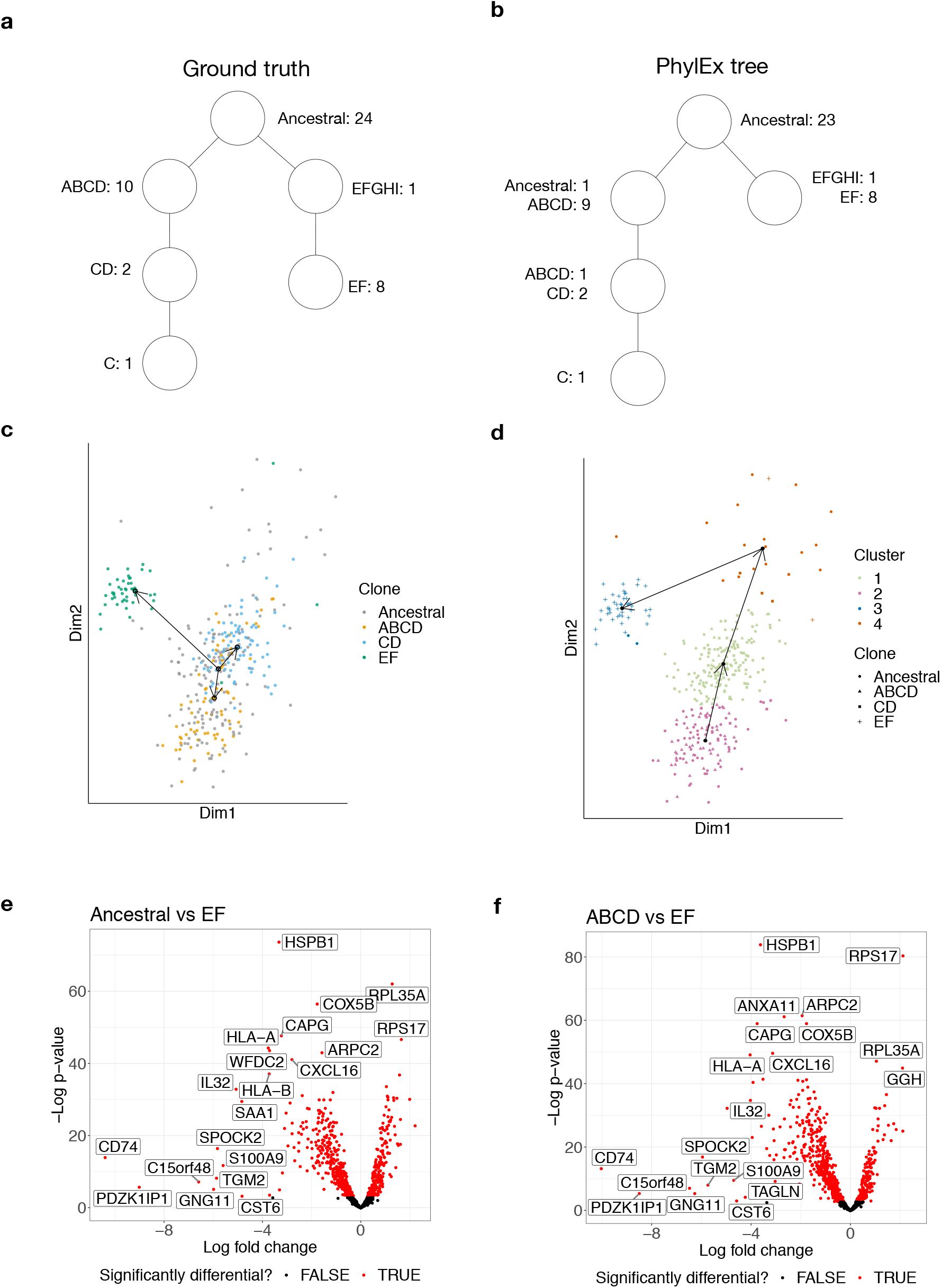
Analysis of HGSOC cell line. a. Ground truth tree with the number of SNVs assigned to each clone indicated beside the clone name. b. The inferred tree from PhylEx with the number of SNVs of the ground truth clone indicated. The plot of the gene expressions for cells on ZINB-WaVE dimensions: c. cells are color-coded after assigning to the clonal-tree output from PhylEx, and the trajectory analysis result is overlayed on the figure with the ancestral clone specified as the starting cluster; d. clustering of cells using *mclust* with the trajectory analysis with starting cluster unspecified. The results of differential gene expression analysis using volcano plots: e. the EF clone to the Ancestral clone, and f. the EF clone to the ABCD clone.

**Table 1:**
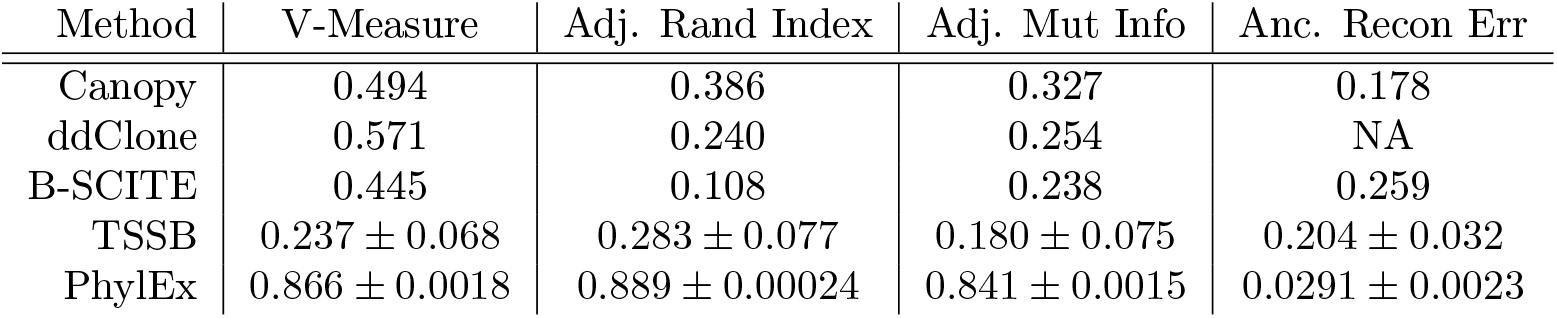
Performance metric comparing PhylEx to Canopy, TSSB, B-SCITE, and ddClone. Used *n* = 20 runs for PhylEx and TSSB. Canopy and B-SCITE were ran with four MCMC chains.

| Method | V-Measure | Adj. Rand Index | Adj. Mut Info | Anc. Recon Err |
| --- | --- | --- | --- | --- |
| Canopy | 0.494 | 0.386 | 0.327 | 0.178 |
| ddClone | 0.571 | 0.240 | 0.254 | NA |
| B-SCITE | 0.445 | 0.108 | 0.238 | 0.259 |
| TSSB | $0.237 \pm 0.068$ | $0.283 \pm 0.077$ | $0.180 \pm 0.075$ | $0.204 \pm 0.032$ |
| PhylEx | $0.866 \pm 0.0018$ | $0.889 \pm 0.00024$ | $0.841 \pm 0.0015$ | $0.0291 \pm 0.0023$ |

### Phylo-phenotypic analysis: PhylEx clones are differentially expressed

To demonstrate PhylEx’s ability to perform phylo-phenotypic analysis we performed gene expression analysis on PhylEx clones based on the HGSOC Smart-Seq3 scRNA-seq data. We cannot evaluate the correctness of cell-to-clone assignment as ground truth does not exist. However, the co-clustering of SNVs and the cells to clones indicates its correctness (Supplementary Figure 7).

We selected the 1,000 genes with the most variable expression pattern for analysis. We used a zero-inflated negative binomial model (ZINB-WaVE) [21] to reduce the dimensionality of the gene expressions data to 2-dimensions. There was a clear separation between the expression of the EF clade (OV2295R) and the primary ABCD clade (OV2295) (Figure 3 **c**). Additionally, cells assigned to CD subclone exhibited a clear separation from the parental ABCD clone (Supplementary Figure 8 **a**). We repeated this analysis using t-SNE [22], another dimensionality reduction technique. A subset of the cells assigned to the ancestral clone, and cells assigned to the EF clone, were well separated (Supplementary Figure 8 **b**). The exhibition of cluster-specific phenotypes, obtained through two independent methods, provides biological evidence of the capacity of PhylEx for phylo-phenotypic analysis. We note that PhylEx only uses SNV allele count data from the scRNA sequencing. As a result PhylEx is blind to gene expression when assigning cells to clones and thus separation of clones in gene expression space is not guaranteed.

We next sought to explore the relationship between pseudo-time trajectories and evolutionary history. Pseudo-time is a popular approach for looking at dynamic changes in gene expression over time. It was first applied in developmental biology studies [23], but is increasingly being used in cancer studies [24]. An open question in the cancer context is whether pseudo-time trajectories reflect evolutionary history. As pseudo-time analysis is based purely on gene expression, this is not guaranteed. We applied the pseudo-time method Slingshot [25] on the 2-dimensional representation obtained by ZINB-WaVE and t-SNE with the cells clustered by PhylEx clones and by gene expression using mclust [26]. Trajectories inferred by slingshot when using PhylEx clusters did not reflect the evolutionary histories. Instead, the parent-child clones ABCD and CD appear as siblings in the ZINB-WaVE dimensions (Figure 3 **c**). The gene expression based clustering using mclust was significantly different from the clone based clusters, and as a result the pseudo-time trajectories differ dramatically from the evolutionary history (Figure 3 **d**). These results suggest that phylo-phenotypic clone based analysis will lead to significantly different interpretations of the data than ones based purely on gene expression.

We performed differential gene expression analysis (DGE) using edgeR [27, 28] to compare the three major clones: (1) the Ancestral clone, the ABCD clone, and the EFclone (Figure 3 E,F). The resulting volcano plots reveal an abundance of differentially expressed genes between the Ancestral/ABCD dominant in the primary tumour, and the EF clone dominant in the relapse. There appears to be immuno-editing in the relapse clone, manifested by a substantial number of down-regulated immune system genes. We performed gene set enrichment analysis (GSEA) using limma package [29] on the set MSigDB C5 (gene ontology) [30]. Several pathways related to the immune system were significantly down-regulated in the EF clone compared to the ABCD clone (Table 2, Supplementary Table 2).

**Table 2:** Gene set enrichment analysis results comparing the ABCD clone to the EF clone. Top 10 most significantly down regulated pathways are shown.

| Gene ontology | P-Value | FDR |
| --- | --- | --- |
| MHC protein complex | 4.85e-15 | 1.32e-11 |
| Antigen processing and presentation of endogenous antigen | 6.40e-15 | 1.32e-11 |
| MHC class I protein complex | 6.57e-14 | 8.08e-11 |
| Response to type I interferon | 2.86e-11 | 1.60e-08 |
| Antigen processing and presentation of endogenous peptide antigen | 2.61e-10 | 1.14e-07 |
| Positive regulation of T cell mediated cytotoxicity | 7.96e-09 | 2.51e-06 |
| Interferon Gamma mediated signaling pathway | 2.08e-08 | 5.72e-06 |
| Regulation of T cell mediated cytotoxicity | 2.37e-08 | 6.07e-06 |
| Positive regulation of antigen processing and presentation | 4.99e-07 | 9.59e-05 |
| Detection of other organism | 6.87e-07 | 1.28e-04 |

Taken together, these results demonstrate that additional insights obtained by comparing expression profiles in the context of PhylEx clones. PhylEx provides the capacity for phylo-phenotypic analysis which can be used to dissect the tumor gene expression patterns beyond what is possible with current single-cell expression analysis tools

### Multi-region HER2+ breast cancer analysis

We generated Smart-Seq3 scRNA-seq and bulk whole-exome DNA sequencing data for five spatially distinct regions of an untreated HER2+ breast cancer tumor (Methods). We applied PhylEx to 369 cells and 418 SNVs that were available after pre-processing. The PhylEx MAP tree was a linear expansion, i.e., a path (Figure 4 **a**), after restriction to non-empty clones (Supplementary Table 3, Supplementary Table 4). The clone fraction appeared to be well-mixed in each region (Figure 4 **d**, Supplementary Table 5). The clone fraction of regions D, E differed from the other regions; this is perhaps explained by the remoteness of these regions to other regions (Supplementary Figure 9).

**Figure 4:**
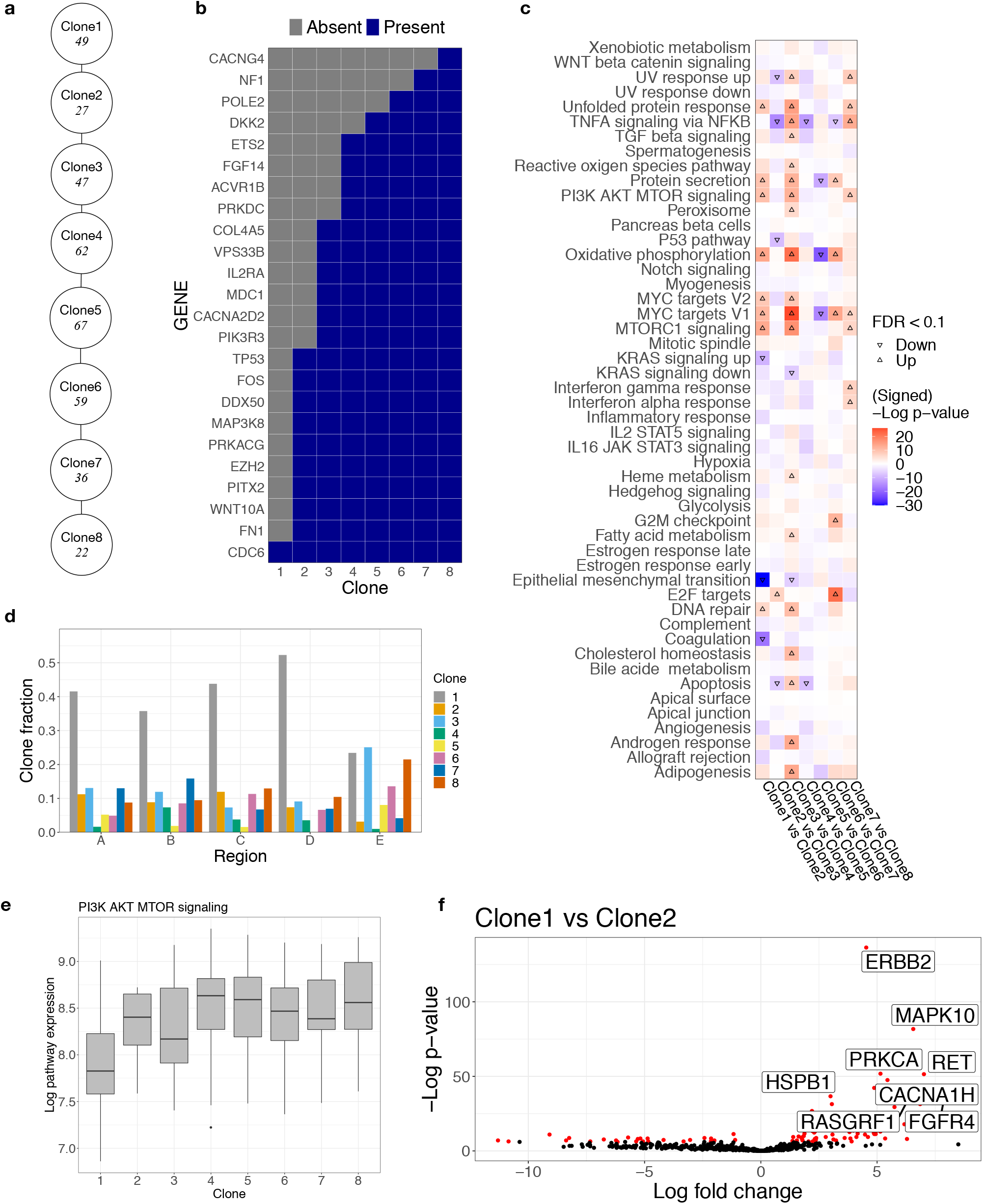
Multi-region HER2+ breast cancer analysis. a. PhylEx inferred tree with the number of cells assigned to each clone shown under the clone label. b. Mutation absence/presence heatmap. c. Heatmap of gene set enrichment analysis on Hallmark pathways to compare parent-child clones. d. Clone (cellular) fraction plot for each clone by region. e. Box-plot of expression levels for PI3K AKT MTOR signaling pathway by clone. f. Differential gene expression analysis to compare progenitor cells assigned to Clone 2 to the cells to Clone 1.

We retrieved the NanoString PanCancer human pathway panel gene list of *n* = 770 curated genes (NanoString Technologies, Seattle, WA) for the downstream analysis. Focusing on this set of genes helps to identify driver mutations for each clone. Among the 432 SNVs used in our analysis, 24 overlapped the NanoString list (Figure 4 **b**). We identified a mutation in CDC6 in the progenitor clone (Clone 1), implicating changes to the cell replication mechanism, and identified a mutation in TP53 and MAP3K8 in Clone 2, hinting at the proliferation of cancer beginning at Clone 2. In Clone 3, we noted mutations to genes involved in PI3K and MAPK pathways (PIK3R3, CACNA2D2) and to MDC1 (DNA repair). Clone 4 appears to be characterized by changes to the RAS pathway as evidenced by mutations to ETS2. Overall, the clonal-tree provides a vital context in which to analyze and inspect mutations in cancer.

We performed gene set enrichment analysis on the MSigDB Hallmark gene sets to compare the parent-child clones (Figure 4 **c**). GSEA revealed a significant increase of PI3K AKT MTOR signaling pathway expression, one of the hallmarks of cancer, in Clone 2 compared to Clone 1. An in-depth inspection of the expression revealed an upregulation of PI3K AKT MTOR signaling pathway in all clones descending from Clone 1 (Figure 4 **e**). We then performed DGE to compare the clones (Figure 4 **f**, Supplementary Figure 10). By performing differential gene expression analysis, we confirmed an overexpression of ERBB2 in Clone 2 compared to Clone 1 (FDR < 0.1). We observed that 49 cells assigned to Clone 1 had a mutation in CDC6 and only two other mutations, perhaps indicating that its cells more closely resemble normal cells than the cancer cells.

Overall, the PhylEx analysis identifies the driver mutations (Figure 4 **b**), elucidates spatial distribution of the clones (Figure 4 **c**), and facilitates a downstream analysis of scRNA expression data that sheds light on the clones’ functional characteristics (Figure 4 **d**-**f**).

## Discussion

In this work we have presented a new method for inferring clonal-trees. By integrating scRNA data during inference we are able to improve reconstruction accuracy. A key benefit of this approach is that we also assign scRNA cells to clones and generate clonal expression profiles. With the prevalence of bulk DNA sequencing and rapidly growing studies conducting scRNA-seq, we expect that PhylEx will prove profitable to cancer researchers studying the functional implications of cancer evolution. We have shown how PhylEx can be used to enhances downstream analysis by providing a clonal-tree and the context to compare the clones’ functional states – revealing the interplay between the evolutionary process and the clones’ phenotypes.

We established that specialized methods for integrating bulk with single-cell transcriptomics are necessary. By modeling read counts PhylEx bypasses the need for a variant calling in the scRNA data and avoids adding uncertainty stemming from dichotomizing counts into binary values.

PhylEx improves over bulk-based clone reconstruction method and should be the preferred choice for inferring the guide tree needed for Cardelino. Similarly, PhylEx is a strong alternative to DLP scDNA-seq for mapping expression profiles to clones using methods such as clonealign. PhylEx, using only a single region bulk sequencing when combined with scRNA-seq, outperforms state-of-the-art bulk-based methods using multi-region bulk data. Even in the settings where replicates are available via multi-region sampling, PhylEx outperforms the bulk-based methods by integrating scRNA-seq.

Another related method is Cardelino-free, which performs *de novo* construction of the clonal-tree when a guide tree is unavailable. However, this amounts to building a tree using only the scRNA-seq data; indeed, the authors of Cardelino found the performance to deteriorate compared to when a guide tree is provided [13]. Building a phylogenetic tree on scRNA-seq has also been considered (e.g., [31]). The phylogenetic tree inferred this way can also serve to guide the downstream analysis as PhylEx – however, we consider these methods to be tangential to PhylEx in the same way that single-cell phylogenies are different from bulk deconvolution and clonal-tree reconstruction problem.

By including the bulk data in the analysis, we can significantly decrease the number of false-positive SNVs detected and incorporate copy number information into the clonal-tree reconstruction. Future extensions of PhylEx to characterize clones by both somatic mutations and copy number profiles have the potential for detecting subclonal copy number information. Inferring subclonal copy numbers is inherently challenging to achieve using only the bulk sequencing data and is only currently feasible using specialized sequencing techniques such as DLP WGS on single cells [32, 14, 33]. PhylEx represents substantial progress in reconstructing the full evolutionary trajectory of cancer.

## Methods

### PhylEx probabilistic model

PhylEx performs a Bayesian posterior inference over the clonal-tree, *T*, and cellular prevalences, ***ϕ***, given the bulk data ***B*** and single-cell data ***S***. The nodes of the tree *T* represent clones, and we will use the term node and clone interchangeably. The likelihood of the bulk and single-cell data is assumed to be conditionally independent given the tree and assignment of SNVs to nodes (i.e., clones) of the tree, ***z***:

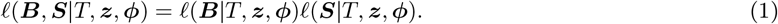

The posterior distribution is expressed in terms of this likelihood and the prior distribution over *T*, ***z, ϕ*** as follows:

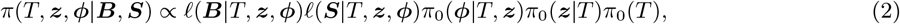

where *π*_0_(***ϕ*** |*T*, ***z***), *π*_0_(***z***|*T*), *π*_0_(*T*) denote the prior distributions over the cellular prevalences, assignment of SNVs to clones, and the tree respectively.

The prior on the clonal-tree is a tree-structured stick-breaking process (TSSB) [15]. TSSB is a Bayesian non-parametric prior defined on infinite trees that learns the tree shape from the data. TSSB has proven to be quite useful in various cancer phylogenetics problems involving both in the context of single-cell and bulk data analysis, e.g., [3, 4, 5].

The prior on the assignment of SNVs to nodes of the tree under TSSB follows a process resembling Polya’s urn scheme but with the hierarchy of urns. Starting from the root node, the prior probability of assigning an SNV to a node *u* is equal to [15]

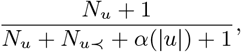

where *N*_*u*_ is the number of SNVs assigned to *u* and *N*_*u≺*_ is the number of SNVs assigned to *u* or its descendants. This indicates that an SNV is assigned to a node with probability that is proportional to the number of SNVs already assigned to *u*, which is the property that enforces clustering of SNVs. We have a function *α*(|*u*|) = *α*_0_*λ*^|*u*|^, which translates to the probability of assigning an SNV to a new node below *u*. This function is governed by hyperparameters *α*_0_ > 0, *λ*∈(0, 1), which decays to 0 as the depth of the node, |*u*|, increases. It serves to ensure that the tree does not grow infinitely. The prior distribution on the cellular prevalences is as specified in [3]. In essense, this amounts to converting the cellular prevalences to clone fraction,

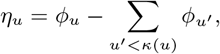

where *κ*(*u*) denotes the set of children nodes of *u*. Note that Σ_*u*_ *η*_*u*_ = 1 for a fixed tree *T* and hence, we can place a Dirichlet distribution on *η*_*u*_ as a prior distribution, conditioned on tree *T*.

We use a likelihood model similar to [2], where clonal copy number information is assumed to be available along with the number of reads mapping to the variant and reference alleles. Copy number analysis is part of a standard bulk data analysis pipeline, which typically includes a normal control sample. We assume clonal major and minor

copy numbers, (*M*_*n*_, *m*_*n*_), covering each SNV, *n* = 1, …, *N*. The bulk data is denoted by 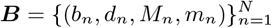, where *b*_*n*_, *d*_*n*_ denote the variant reads and depth at locus *n*. We assume site independence conditional on *T*, ***z***:

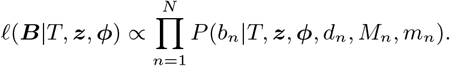

All possible copy number profiles is marginalized to compute the likelihood of the observed reads

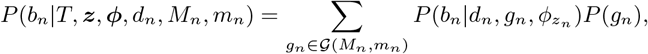

where *𝒢*(*M*_*n*_, *m*_*n*_) = *{A, B, AA, AB, BB, AAA*, …*}*are possible genotypes compatible with a given major and minor copy number profile. The probability distribution for the variant reads is given by Binomial distribution:

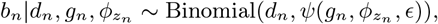

with 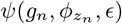 being the probability of success given as a function of the genotype, cellular prevalence of clone *z*_*n*_, and sequencing error probability, *ϵ*. Letting *v*(*g*), *c*(*g*) be the number of variant alleles and total copy numbers for a genotype *g*, the success probability is given by,

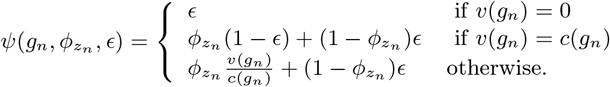

The scRNA-seq data is denoted by 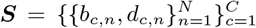, where *C* denotes the number of cells. We assume that the scRNA-seq likelihood is conditionally independent over cell and locus given *T*, ***z*** and cell-to-clone membership, 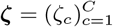:

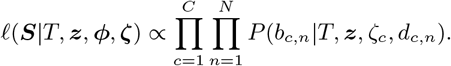

The cell-to-clone membership variable completely determines the SNVs harbored by cells. For a cell assigned to node *u*, it inherits all of the SNVs assigned to ancestral nodes of *u* in *T*. We denote the mutation status of cell *c* for locus *n* by *µ*_*c,n*_ ∈*{*0, 1*}*, which can be seen as a function of *T*, ***z***, *ζ*_*c*_.

We model the number of variant reads, *b*_*c,n*_, for each cell *c* and locus *n* using a mixture of two Beta-Binomial distributions. The mixture is necessary to account for sparse nature of scRNA-seq, in particular, we need one distribution to model monoallelic expression (i.e., dropout) and another for biallelic distribution (see e.g., [31]).

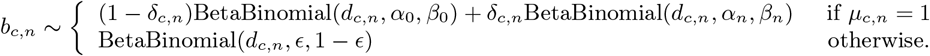

where *α*_0_, *β*_0_, *α*_*n*_, *β*_*n*_ are hyperparameters of the model (Supplement). For the monoallelic distribution, we choose the hyperparameters *α*_0_, *β*_0_ to be small, e.g., *α*_0_ = *β*_0_ *<* 0.05 (Supplementary Figure 11). This ensures that most of the mass is centered at near 0 and *d*_*c,n*_. Such distribution guards against cases where a cell harbours the mutation but only the reference allele is expressed due to the stochastic nature of the transcriptional process, leading to unbalanced expression of alleles in scRNA-seq; similar techniques are employed in [34, 31, 13]. The hyper-parameters *α*_*n*_, *β*_*n*_ for each site *j* are estimated as part of data pre-processing step (Supplement). In the computation of the likelihood, we marginalize out *δ*_*c,n*_.

The prior probability of cell assignment to clone *u* can be given by the clone fraction, *η*_*u*_. However, such an assumption may not hold as cells with certain characteristics may be preferentially selected for sequencing. Therefore, we use Uniform distribution:

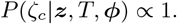

In evaluating the single-cell component of the likelihood for a given tree, we marginalize over the cell-to-clone assignments,

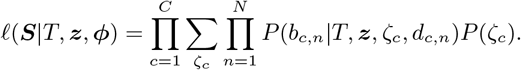

### Approval and collection of clinical material for breast cancer samples

Fresh primary tumor resections were obtained from breast cancer patients at Karolinska University Hospital and Stockholm South General Hospital. Experimental procedures and protocols were approved by the regional ethics review board (Etikprövningsnämnden) in Stockholm, with reference numbers 2016/957-31 and 2017/742-32. Biobank approval was obtained from the Stockholm medical biobank. Before surgery, informed consent in accordance with the Declaration of Helsinki was given to patients for signature.

### Whole-exome sequencing for breast cancer samples

Tumor resections and matching dermal biopsies from 4 individual breast cancer patients were freshly collected. Tissues were manually homogenized and genomic DNA samples were isolated by using the QIAamp DNA mini kit (QIAGEN). The library was prepared by using Twist Bioscience Human Core Exome kit (Twist Bioscience) according to the manufacture protocol. The bulk DNA samples were then sequenced in a S4 flow cell lane by the NovaSeq 6000 platform (Illumina) at the National Genomics Infrastructure, Science for Life Laboratory, Uppsala.

### Breast cancer sample preparation for Smart-Seq3 method

Tissues were homogenized and cells were released by using the gentleMACS™ Octo Dissociator with Heaters and the human tumor dissociation kit (both from Miltenyi Biotec), according to the manufacturer protocols. Afterwards, the cells were washed two times with F12-DMEM medium (Gibco) and collected by centrifugation at 300g for 5 minutes. The single cell suspensions were further generated by passing the re-suspended cells through the 70 mm cell strainers. The single cell suspensions were then further stained with the Zombie Aqua Fixable viability dye (Biolegend, 423101) at room temperature for 20 minutes, then washed with Phosphate Buffered Saline (PBS). The cells were incubated with Human TruStain Fc block (Biolegend, 422302) for 10 minutes to limit unspecific antibody binding, then stained for 20 minutes with anti-EPCAM (1:40, Biolegend, 324206) and anti-CD45 (1:40, Biolegend, 304021) in FACS buffer (PBS + 0.5 % Bovine Serum Albumin). The cells were subsequently washed and resuspended in FACS buffer. Fluorescence-activated cell sorting (FACS) using an influx flow cytometer (BD Biosciences) was performed to sort live EPCAM+CD45-single cells into 384 well plates for Smart-Seq3 analysis. Single stain controls (cells and beads) and fluorescence minus one controls (FMO), containing all the fluorochromes in the panel except the one being measured, were used to set voltages and to define the proper gating strategy. Equal numbers of cells from each tumor region was sorted onto the same plate and two plates were prepared in total.

### Ovarian cancer cell lines preparation for Smart-Seq3

Culture of ovarian cancer cell lines OV2295, TOV2295 and OV2295R cells were cultured in a 1:1 mix of Media 199 (Sigma Aldrich) and MCDB 105 (Sigma Aldrich) supplemented with 10% FBS in a humidified environment at 37C.

or single cell RNA sequencing, all cells used in this study were sorted on a BD Influx into 384 well plates using index-sorting and single-cell purity mode directly into lysis buffer(6,67% Polyethylene Glycol, 0.1% Triton X-100, RNAse Inhibitor (Takara), dNTPs (0.67mM/each) and Oligo-dT (0.67uM)). Sorted plates were stored at-80C and thawed immediately prior to library generation.

### Smart-Seq3 library preparation and sequencing

For single cell RNAseq libraries, the Smart-Seq3 method was used according to the published protocol (PMID: 32518404). In brief, plates were quickly centrifuged before reverse transcription (25mM Tris-HCl pH 8.3 (Sigma), 30mM NaCl (ThermoFisher), 2,5mM MgCl2 (ThermoFisher), 1mM GTP (ThermoFisher), 8mM DTT (ThermoFisher), 0,5u/ul RNase inhibitor (Takara), 2uM TSO (IDT), 2u/ul Maxima H-minus reverse transcriptase (ThermoFisher)) and amplified using KAPA HiFI Hotstart polymerase (Roche) to generate full length cDNA libraries (22 cycles PCR). Final library concentrations were determined and normalized for each cell using Picogreen. Diluted cDNA of 100pg per sample was used for tagmentation (Nextera Library Preparation Kit, Illumina, ATM at 0,1uL per cell). The final samples were analyzed using a Bioanalyzer (Hi-Sensitivity Kit, Agilent) and sent for sequencing on a Novaseq S Prime lane, PE 2×150bp (Illumina). Library quality was compared to index sorting results to confirm that negative wells yielded low complexity libraries.

### Processing of Smart-Seq3 sequencing files

Individual fastq files for the cells are obtained using Illumina bcl2fq tool, then converted to ubam format with cell and UMI tags using a script that detect Smart-Seq 3 specific pattern at the beginning of reads with UMI. STAR version 7.1 with GRCH37 version of Human Genome and Ensembl version 75 annotations were used to align the reads. UMI-tools[35] was then used to correct the UMI and group UMI reads. An in-house script was used to intersect the reads with bam files and obtain read and UMI counts for each gene.

### Mutation calling from exome sequencing results

Mutations were called using Mutect2[36] software in multi-sample mode, with recommended filters. Point mutations that passed the filters were selected for downstream analysis.

## Code and data availability

Code implementing PhylEx and the R code for analysis is available on Github at https://github.com/junseonghwan/PhylExAnalysis. The simulation data and results, processed bulk DNA-seq and scRNA-seq data for HGSOC and HER2+ data along with the results are available at https://doi.org/10.5281/zenodo.4533670.

## Acknowledgements

The computations were enabled by resources provided by the Swedish National Infrastructure for Computing (SNIC) at Uppsala University and Linköping University partially funded by the Swedish Research Council through grant agreement no. 2018-05973. This project was made possible through funding by the Michael Smith Foundation for Health Research Scholar Award [18245 to AR] and by generous support from the Swedish Foundation for Strategic Research grant BD15-0043. The first author was supported by Postdoctoral Fellowship from the Natural Sciences and Engineering Research Council of Canada.

## Footnotes

1 TOV2295R is difficult to grow and we could not use it.

## References

[1] P C Nowell. The clonal evolution of tumor cell populations. Science, 194(4260):23–28, October 1976.

[2] Andrew Roth, Jaswinder Khattra, Damian Yap, Adrian Wan, Emma Laks, Justina Biele, Gavin Ha, Samuel Aparicio, Alexandre Bouchard-Côté, and Sohrab P Shah. PyClone: statistical inference of clonal population structure in cancer. Nature methods, 11(4):396, 2014.

[3] Wei Jiao, Shankar Vembu, Amit G Deshwar, Lincoln Stein, and Quaid Morris. Inferring clonal evolution of tumors from single nucleotide somatic mutations. BMC bioinformatics, 15(1):35, 2014.

[4] Amit G Deshwar, Shankar Vembu, Christina K Yung, Gun Ho Jang, Lincoln Stein, and Quaid Morris. PhyloWGS: reconstructing subclonal composition and evolution from whole-genome sequencing of tumors. Genome biology, 16(1):35, 2015.

[5] Ke Yuan, Thomas Sakoparnig, Florian Markowetz, and Niko Beerenwinkel. Bitphylogeny: a probabilistic framework for reconstructing intra-tumor phylogenies. Genome biology, 16(1):36, 2015.

[6] Yuchao Jiang, Yu Qiu, Andy J Minn, and Nancy R Zhang. Assessing intratumor heterogeneity and tracking longitudinal and spatial clonal evolutionary history by next-generation sequencing. Proceedings of the National Academy of Sciences, 113(37):E5528–E5537, 2016.

[7] Andrew Roth, Andrew McPherson, Emma Laks, Justina Biele, Damian Yap, Adrian Wan, Maia A Smith, Cydney B Nielsen, Jessica N McAlpine, Samuel Aparicio, Alexandre Bouchard-Côté, and Sohrab P Shah. Clonal genotype and population structure inference from single-cell tumor sequencing. Nat. Methods, 13(7):573–576, July 2016.

[8] Katharina Jahn, Jack Kuipers, and Niko Beerenwinkel. Tree inference for single-cell data. Genome biology, 17(1):86, 2016.

[9] Edith M Ross and Florian Markowetz. OncoNEM: inferring tumor evolution from single-cell sequencing data. Genome biology, 17(1):1–14, 2016.

[10] Sohrab Salehi, Adi Steif, Andrew Roth, Samuel Aparicio, Alexandre Bouchard-Côté, and Sohrab P Shah. ddclone: joint statistical inference of clonal populations from single cell and bulk tumour sequencing data. Genome biology, 18(1):44, 2017.

[11] Salem Malikic, Katharina Jahn, Jack Kuipers, Cenk Sahinalp, and Niko Beerenwinkel. Integrative inference of subclonal tumour evolution from single-cell and bulk sequencing data. bioRxiv, page 234914, 2017.

[12] Kieran R Campbell, Adi Steif, Emma Laks, Hans Zahn, Daniel Lai, Andrew McPherson, Hossein Farahani, Farhia Kabeer, Ciara O’Flanagan, Justina Biele, Jazmine Brimhall, Beixi Wang, Pascale Walters, IMAXT Consortium, Alexandre Bouchard-Côté, Samuel Aparicio, and Sohrab P Shah. clonealign: statistical integration of independent single-cell RNA and DNA sequencing data from human cancers. Genome Biol., 20(1):54, March 2019.

[13] Davis J McCarthy, Raghd Rostom, Yuanhua Huang, Daniel J Kunz, Petr Danecek, Marc Jan Bonder, Tzachi Hagai, Ruqian Lyu, Wenyi Wang, Daniel J Gaffney, et al. Cardelino: computational integration of somatic clonal substructure and single-cell transcriptomes. Nature Methods, 17(4):414–421, 2020.

[14] Emma Laks, Andrew McPherson, Hans Zahn, Daniel Lai, Adi Steif, Jazmine Brimhall, Justina Biele, Beixi Wang, Tehmina Masud, Jerome Ting, Diljot Grewal, Cydney Nielsen, Samantha Leung, Viktoria Bojilova, Maia Smith, Oleg Golovko, Steven Poon, Peter Eirew, Farhia Kabeer, Teresa Ruiz de Algara, So Ra Lee, M Jafar Taghiyar, Curtis Huebner, Jessica Ngo, Tim Chan, Spencer Vatrt-Watts, Pascale Walters, Nafis Abrar, Sophia Chan, Matt Wiens, Lauren Martin, R Wilder Scott, T Michael Underhill, Elizabeth Chavez, Christian Steidl, Daniel Da Costa, Yussanne Ma, Robin J N Coope, Richard Corbett, Stephen Pleasance, Richard Moore, Andrew J Mungall, Colin Mar, Fergus Cafferty, Karen Gelmon, Stephen Chia, CRUK IMAXT Grand Challenge Team, Marco A Marra, Carl Hansen, Sohrab P Shah, and Samuel Aparicio. Clonal decomposition and DNA replication states defined by scaled Single-Cell genome sequencing. Cell, 179(5):1207–1221.e22, November 2019.

[15] Ryan P Adams, Zoubin Ghahramani, and Michael I Jordan. Tree-structured stick breaking for hierarchical data. In Advances in Neural Information Processing Systems, pages 19–27, 2010.

[16] Radford M Neal. Slice sampling. Annals of statistics, pages 705–741, 2003.

[17] Anton J M Larsson, Per Johnsson, Michael Hagemann-Jensen, Leonard Hartmanis, Omid R Faridani, Björn Reinius, Åsa Segerstolpe, Chloe M Rivera, Bing Ren, and Rickard Sandberg. Genomic encoding of transcriptional burst kinetics. Nature, 565(7738):251–254, January 2019.

[18] Serena Nik-Zainal, Peter Van Loo, David C Wedge, Ludmil B Alexandrov, Christopher D Greenman, King Wai Lau, Keiran Raine, David Jones, John Marshall, Manasa Ramakrishna, et al. The life history of 21 breast cancers. Cell, 149(5):994–1007, 2012.

[19] Russell Schwartz and Alejandro A Schäffer. The evolution of tumour phylogenetics: principles and practice. Nature Reviews Genetics, 18(4):213–229, 2017.

[20] Michael Hagemann-Jensen, Christoph Ziegenhain, Ping Chen, Daniel Ramsköld, Gert-Jan Hendriks, Anton JM Larsson, Omid R Faridani, and Rickard Sandberg. Single-cell RNA counting at allele and isoform resolution using Smart-Seq3. Nature Biotechnology, 38(6):708–714, 2020.

[21] Davide Risso, Fanny Perraudeau, Svetlana Gribkova, Sandrine Dudoit, and Jean-Philippe Vert. A general and flexible method for signal extraction from single-cell RNA-seq data. Nature communications, 9(1):1–17, 2018.

[22] Laurens van der Maaten and Geoffrey Hinton. Visualizing data using t-SNE. Journal of machine learning research, 9(Nov):2579–2605, 2008.

[23] Cole Trapnell, Davide Cacchiarelli, Jonna Grimsby, Prapti Pokharel, Shuqiang Li, Michael Morse, Niall J Lennon, Kenneth J Livak, Tarjei S Mikkelsen, and John L Rinn. Pseudo-temporal ordering of individual cells reveals dynamics and regulators of cell fate decisions. Nature biotechnology, 32(4):381, 2014.

[24] Jean Fan, Kamil Slowikowski, and Fan Zhang. Single-cell transcriptomics in cancer: Computational challenges and opportunities. Experimental & Molecular Medicine, 52(9):1452–1465, 2020.

[25] Kelly Street, Davide Risso, Russell B Fletcher, Diya Das, John Ngai, Nir Yosef, Elizabeth Purdom, and Sandrine Dudoit. Slingshot: cell lineage and pseudotime inference for single-cell transcriptomics. BMC genomics, 19(1):477, 2018.

[26] Luca Scrucca, Michael Fop, T. Brendan Murphy, and Adrian E. Raftery. mclust 5: clustering, classification and density estimation using Gaussian finite mixture models. The R Journal, 8(1):289–317, 2016.

[27] Mark D Robinson, Davis J McCarthy, and Gordon K Smyth. edgeR: a bioconductor package for differential expression analysis of digital gene expression data. Bioinformatics, 26(1):139–140, January 2010.

[28] Davis J McCarthy, Yunshun Chen, and Gordon K Smyth. Differential expression analysis of multifactor RNA-Seq experiments with respect to biological variation. Nucleic Acids Res., 40(10):4288–4297, May 2012.

[29] Matthew E Ritchie, Belinda Phipson, D. Wu, Yifang Hu, Charity W Law, Wei Shi, and Gordon K Smyth. limma powers differential expression analyses for RNA-sequencing and microarray studies. Nucleic Acids Res., 43(7):e47, April 2015.

[30] Arthur Liberzon, Aravind Subramanian, Reid Pinchback, Helga Thorvaldsdóttir, Pablo Tamayo, and Jill P Mesirov. Molecular signatures database (MSigDB) 3.0. Bioinformatics, 27(12):1739–1740, 2011.

[31] Zilu Zhou, Bihui Xu, Andy Minn, and Nancy R Zhang. DENDRO: genetic heterogeneity profiling and subclone detection by single-cell RNA sequencing. Genome Biology, 21(1):1–15, 2020.

[32] Hans Zahn, Adi Steif, Emma Laks, Peter Eirew, Michael VanInsberghe, Sohrab P Shah, Samuel Aparicio, and Carl L Hansen. Scalable whole-genome single-cell library preparation without preamplification. Nature methods, 14(2):167, 2017.

[33] Fatemeh Dorri, Sohrab Salehi, Kevin Chern, Tyler Funnell, Marc Williams, Daniel Lai, Mirela Andronescu, Kieran R Campbell, Andrew McPherson, Samuel Aparicio, Andrew Roth, Sohrab Shah, and Alexandre Bouchard-Côté. Efficient Bayesian inference of phylogenetic trees from large scale, low-depth genome-wide single-cell data. bioRxiv, 2020.

[34] Yuchao Jiang, Nancy R Zhang, and Mingyao Li. SCALE: modeling allele-specific gene expression by single-cell RNA sequencing. Genome biology, 18(1):74, 2017.

[35] Tom Smith, Andreas Heger, and Ian Sudbery. UMI-tools: modeling sequencing errors in unique molecular identifiers to improve quantification accuracy. Genome research, 27(3):491–499, 2017.

[36] David Benjamin, Takuto Sato, Kristian Cibulskis, Gad Getz, Chip Stewart, and Lee Lichtenstein. Calling somatic SNVs and Indels with Mutect2. bioRxiv, 2019.

