## Supplementary Materials for "PhylEx: Accurate reconstruction of clonal structure via integrated analysis of bulk DNA-seq and single cell RNA-seq data"

### Method details

#### Input data preparation

The bulk tumor and matching normal samples are to be pre-processed following standard guidelines as per GATK standard practice [1]. We recommend to use Strelka or Mutect2 to detect SNVs, their position in the genome (loci), and finally, the variant and reference alleles for each loci [2, 3]. The PASS filter can be used to select the high-confidence SNVs. PhylEx requires major and minor copy number profiles for the SNVs, which can be obtained from the bulk samples using TitanCNA [4] or other similar softwares. The scRNA-seq can be aligned using STAR aligner [5]. From the algined scRNA-seq, the reads mapping to the variant and the reference can be read off at the loci and the alleles identified from the bulk DNA-seq. For this purpose we used an Rsamtools package [6].

#### High-grade serous ovarian cancer cell-line evaluation

The pseudobulk DNA-seq data from tumor and matching normal samples are pre-processed as above. The copy number profiles are obtained from the bulk samples using TitanCNA and the scRNA-seq is aligned using STAR aligner. A list of SNVs are provided in the data repository provided in [7]; therefore, we did not need to make variant calls. Among the list of SNVs provided in [7], 634 SNVs were found to be exonic. We further filtered these set of SNVs using scRNA-seq data. For each loci $n = 1, ..., 634$, we selected it for analysis if there were at least two cells such that $b_{c,n} >= 2$, which resulted in 67 SNVs. Including SNVs that does not have sufficient single-cell coverage does not help to evaluate PhylEx's capacity for inferring the branching events and the ancestral relationship. We retrieved the reads at each of these 67 locis for each cell using *Rsamtools*.

### Evaluation metrics

We used V-measure, adjusted rand index, and adjusted mutual information as implemented in scikit-learn (version 0.23.1) [8]. To evaluate the reconstruction accuracy, we defined an ancestral reconstruction error, defined on a pair of SNVs. For two nodes $u, v$ in the tree, we say $u < v$ to mean that $u$ is ancestral to $v$. We can extend this definition to SNVs $i, j$. We will say $i < j$ if and only if $i$ is assigned to node $u$ and $j$ to $v$ such that $u < v$. We formulate an ancestral matrix of dimension $N \times N$, where the $(i, j)$-th is set to 1 if $i < j$. We denote the ancestral matrix for the ground truth SNV-to-clone assignment by $A^*$, then we can compute the absolute error (AE) of an ancestral matrix $A$ by

$$\mathrm{AE}(A^*, A) = \sum_{i,j} |A^*_{i,j} - A_{i,j}|.$$

The mean aboslute error is given by dividing AE by the number of unique pairs.

### PhylEx model

#### Modeling scRNA-seq data

We plotted the histogram of the ratio of variant reads to depth over all sites and cells i.e., $(b_{c,n}/d_{c,n})$ for HGSOC data (Supplementary Figure 11 **a**). This plot clearly depicts the monoallelic nature of the expression data. We plotted the bi-allelic sites by selecting a subset of the data such that $b_{c,n} > 0$, $b_{c,n}/d_{c,n} < 1$ (Supplementary Figure 11 **b**). We have also made similar plots on HER2+ scRNA-seq data (Supplementary Figure 11 **c**, **d**). These plots demonstrate that a mixture distribution is suitable for modeling the read counts from scRNA-seq.

For cell $c$ that harbors the mutation $n$, we assume the following generative process for the scRNA-seq data:

$$\begin{aligned} \delta_{c,n} &\sim \mathrm{Bernoulli}(\delta_0) \\ \chi_{c,n} | \delta_{c,n} &\sim \delta_{c,n} \mathrm{Beta}(\alpha_n, \beta_n) + (1 - \delta_{c,n}) \mathrm{Beta}(\alpha_0, \beta_0) \\ b_{c,n} | d_{c,n}, \chi_{c,n} &\sim \mathrm{Binomial}(d_{c,n}, \chi_{c,n}) \end{aligned} \tag{1}$$

where $\delta_0$ is the probability of bursty expression and $\chi_{c,n}$ is the probability of expression the variant allele. For small values of $\alpha_0, \beta_0$, the monoallelic distribution has most of the mass at the two ends as shown in Supplementary Figure 12 **a-c**. We use $\alpha_0 = \beta_0 = 0.01$ for HGSOC and HER2+ analysis, as most of the mass is placed at the two extremes (Supplementary Figure 11 **a**, **c**).

Our primary interest is in estimating $\alpha_n, \beta_n$ as a part of the preprocessing step. We describe a simple approach that we used in the data analysis. Our approach is to predict $\delta_{c,n}$ first, then estimate $\alpha_n, \beta_n$ using Beta-Binomial conjugacy.

To predict $\delta_{c,n}$, we compute:

$$P(\delta_{c,n} = 1 | b_{c,n}, \alpha^0_n, \beta^0_n, \alpha_0, \beta_0, \delta_0) \propto P(b_{c,n} | \delta_{c,n} = 1, \alpha^0_n, \beta^0_n, \alpha_0, \beta_0, ) P(\delta_{c,n} = 1 | \delta_0),$$

where $\alpha^0_n, \beta^0_n$ quantify initial belief over the parameters for bi-allelic Beta distribution. We set $\alpha^0_n = \beta^0_n = 1$, which defines the Uniform distribution on $[0, 1]$, $\delta_0 = 0.5$, and $\alpha_0 = \beta_0 = 0.05$. Given $\delta_{c,n}$, the hyper parameter update is,

$$\begin{aligned} \alpha_n &= \alpha^0_n + \sum_{n:\delta_{c,n}=1} b_{c,n} \\ \beta_n &= \beta^0_n + \sum_{n:\delta_{c,n}=1} (d_{c,n} - b_{c,n}), \end{aligned}$$

which is the standard update formula for Beta-Binomial hyperparameters.

### Simulated Data Generation

#### Simulating Clone Tree

We generate a binary tree with the root node representing the non-malignant ancestor. The root node has exactly one child, representing progenitor cancer clone with cellular prevalence of 0.5. We grow the tree starting from the progenitor clone such that each node has exactly two children; each child breaks half of the parent's remaining clone fraction: $\phi_u = 0.5\eta_{\rho(u)}$. We stop expanding the tree at a node when the depth reaches the maximum specified or the cellular prevalence of the node falls below minimum threshold. In the simulated experiments, we set the maximum depth to 3 (depth of the root node is 0) and minimum cellular prevalence to 0.05. This results in a binary tree with eight nodes as shown in Supplementary Figure 13 **a**. We have an implementation that allows the cellular prevalence to be randomly sampled but generating the cellular prevalence in a determined manner creates for an interesting and challenging scenario. In particular, there are two pairs of clones that have the same cellular prevalence of 0.125 and 0.0625. Having clones with the same cellular fraction makes it difficult for reconstruction based on variant allele frequencies alone. To simulate the multifurcating tree, we again set the root node to be the healthy clone and it has exactly one child that represents the progenitor cancer clone. Starting from the progenitor cancer clone, we grow the tree by randomly selecting number of children from $\{1, 2, 3, 4\}$. The clone fraction of the children nodes are determined in the same way as for the binary tree. We stop growing the tree when the maximum depth of 3 or the minimum cellular prevalence of 0.05 is reached. Examples of trees generated for simulation studies is shown in Supplementary Figure 13 **b**, **c**.

#### Bulk data generation

To model cancer's complex structural variation and to study the effect of copy number misspecification, we use birth-death process to simulate copy number profiles. Birth-death process is parameterized by the birth rate and the death rate, with maximum copy number of 10 and the absorbing copy number of 0, i.e., once the copy number reaches 0, it does not evolve. We first construct two rate matrices, $\mathbb{Q}_0$ with minimum copy number state of 0 and $\mathbb{Q}_1$ with the minimum copy number state of 1 [9]. The transition matrices are obtained via matrix exponentiation, $\mathbb{P}_0 = \exp(\mathbb{Q}_0)$ and $\mathbb{P}_1 = \exp(\mathbb{Q}_1)$.

Let SNV $n$ be assigned to node $u$ of the tree. We initialize the copy number at the root node with the value of 2. We then separate copy number evolution into three parts. The first part is along the branches from the root node to $u$. We evolve the copy number using $\mathbb{P}_1$, this ensures that there is at least one copy by the time we get to node $u$. Let $X'_{n,u}$ be the copy number at node $u$, we then sample $Y'_{n,u} \sim \text{Binomial}(X'_{n,u} - 1, \xi)$, where $\xi \in (0, 1)$. Then, we set the variant copy profile at $u$ as $Y_{n,u} = Y'_{n,u} + 1$ and set $X_{n,u} = X'_{n,u} - Y'_{n,u}$. The second part is copy number evolution starting at node $u$, which is initialized with copy number profile of $X_{n,u}, Y_{n,u}$. We evolve $X_{n,u}, Y_{n,u}$ independently using $\mathbb{P}_0$. The third part is copy number evolution over all of the other branches, that is, the branches not in the path from root node to $u$ and not in the subtree rooted at $u$. The copy numbers are evolved using $\mathbb{P}_0$ for these branches.

Once we have the full copy number profile at each clone, then we take the weighted averages,

$$\bar{X}_n = \sum_{v \in T} \eta_v X_{n,v} \tag{2}$$

$$\bar{Y}_n = \sum_{v \in T} \eta_v Y_{n,v} \tag{3}$$

$$\bar{D}_n = \sum_{v \in T} \eta_v (X_{n,v} + Y_{n,v}) \tag{4}$$

We round $\bar{X}_n, \bar{Y}_n$, then sort $(\bar{X}_n, \bar{Y}_n)$ to convert it to integer-valued major and minor copy numbers for PhylEx and other softwares used in the study. Note that these copy numbers provided as input to PhylEx does not fully capture the true copy number state of the cancer, which is as we desired.

The bulk data is generated as follows:

$$d_n \sim \text{Poisson}(d_0 \cdot \bar{D}_n/2)$$
$$b_n \sim \text{Binomial}(d_n, \xi_n),$$

where $d_0$ is the desired mean depth, set to $1,000$ in the simulation studies and $\xi_n = \bar{Y}_n/\bar{D}_n$, denoting the probability of observing a variant read. The division by factor of 2 arises when we consider $\bar{D}_n = 2$, e.g., when there is no copy number variation. In that case, to ensure the realized depth has mean $d_0$, we need to divide by 2.

##### Simulating scRNA-seq data

Given the tree and the SNV-to-clone assignment, we first sample a cell-to-clone assignment for each of the cells. Assigning cell to a clone determines its genotype, call it $\mathcal{G}_c$. We randomly select a subset of SNVs to be expressed, $\mathcal{E}_c$. For $n \in \mathcal{G}_c \bigcap \mathcal{E}_c$, we first sample the depth, $d_{c,n}$ from Poisson distribution with mean expression level $e_0$. Then, we do a coin-flip to set $\delta_{c,n}$. If $\delta_{c,n} = 1$, we sample $b_{c,n}$ from Beta-Binomial$(d_{c,n}, \alpha_n, \beta_n)$. The hyperparameters are sampled from uniform distribution with over $(0, max)$ with $max = 10$. If $\delta_{c,n} = 0$, we sample $b_{c,n}$ from Beta-Binomial$(d_{c,n}, \alpha_0, \beta_0)$ with parameters $\alpha_0 = \beta_0 = 0.01$. For $n \in \mathcal{E}_c \setminus \mathcal{G}_c$, we sample $b_{c,n}$ from Beta-Binomial$(d_{c,n}, \epsilon, 1 - \epsilon)$, where $\epsilon = 0.01$ denotes the sequencing error (Supplementary Figure 12 **d**). As the loci $n$ is expressed but the cell does not harbor the SNV, we expect to observe a variant read only in error. For $n \notin \mathcal{E}_c$, we set $d_{c,n} = b_{c,n} = 0$.

#### References

[1] Ryan Poplin, Valentin Ruano-Rubio, Mark A DePristo, Tim J Fennell, Mauricio O Carneiro, Geraldine A Van der Auwera, David E Kling, Laura D Gauthier, Ami Levy-Moonshine, David Roazen, Khalid Shakir, Joel Thibault, Sheila Chandran, Chris Whelan, Monkol Lek, Stacey Gabriel, Mark J Daly, Ben Neale, Daniel G MacArthur, and Eric Banks. Scaling accurate genetic variant discovery to tens of thousands of samples. July 2018.

[2] Christopher T Saunders, Wendy S W Wong, Sajani Swamy, Jennifer Becq, Lisa J Murray, and R Keira Cheetham. Strelka: accurate somatic small-variant calling from sequenced tumor-normal sample pairs. *Bioinformatics*, 28(14):1811–1817, July 2012.

[3] David Benjamin, Takuto Sato, Kristian Cibulskis, Gad Getz, Chip Stewart, and Lee Lichtenstein. Calling somatic SNVs and Indels with Mutect2. *bioRxiv*, 2019.

[4] Gavin Ha, Andrew Roth, Jaswinder Khattra, Julie Ho, Damian Yap, Leah M Prentice, Nataliya Melnyk, Andrew McPherson, Ali Bashashati, Emma Laks, Justina Biele, Jiarui Ding, Alan Le, Jamie Rosner, Karey Shumansky, Marco A Marra, C Blake Gilks, David G Huntsman, Jessica N McAlpine, Samuel Aparicio, and Sohrab P Shah. TITAN: inference of copy number architectures in clonal cell populations from tumor whole-genome sequence data. *Genome Res.*, 24(11):1881–1893, November 2014.

[5] Alexander Dobin, Carrie A Davis, Felix Schlesinger, Jorg Drenkow, Chris Zaleski, Sonali Jha, Philippe Batut, Mark Chaisson, and Thomas R Gingeras. STAR: ultrafast universal RNA-seq aligner. *Bioinformatics*, 29(1):15–21, January 2013.

[6] Martin Morgan, Hervé Pagès, Valerie Obenchain, and Nathaniel Hayden. *Rsamtools: Binary alignment (BAM), FASTA, variant call (BCF), and tabix file import*, 2020. R package version 2.2.3.

[7] Emma Laks, Andrew McPherson, Hans Zahn, Daniel Lai, Adi Steif, Jazmine Brimhall, Justina Biele, Beixi Wang, Tehmina Masud, Jerome Ting, Diljot Grewal, Cydney Nielsen, Samantha Leung, Viktoria Bojilova, Maia Smith, Oleg Golovko, Steven Poon, Peter Eirew, Farhia Kabeer,

Teresa Ruiz de Algara, So Ra Lee, M Jafar Taghiyar, Curtis Huebner, Jessica Ngo, Tim Chan, Spencer Vatrt-Watts, Pascale Walters, Nafis Abrar, Sophia Chan, Matt Wiens, Lauren Martin, R Wilder Scott, T Michael Underhill, Elizabeth Chavez, Christian Steidl, Daniel Da Costa, Yussanne Ma, Robin J N Coope, Richard Corbett, Stephen Pleasance, Richard Moore, Andrew J Mungall, Colin Mar, Fergus Cafferty, Karen Gelmon, Stephen Chia, CRUK IMAXT Grand Challenge Team, Marco A Marra, Carl Hansen, Sohrab P Shah, and Samuel Aparicio. Clonal decomposition and DNA replication states defined by scaled Single-Cell genome sequencing. *Cell*, 179(5):1207–1221.e22, November 2019.

[8] F. Pedregosa, G. Varoquaux, A. Gramfort, V. Michel, B. Thirion, O. Grisel, M. Blondel, P. Prettenhofer, R. Weiss, V. Dubourg, J. Vanderplas, A. Passos, D. Cournapeau, M. Brucher, M. Perrot, and E. Duchesnay. Scikit-learn: Machine learning in Python. *Journal of Machine Learning Research*, 12:2825–2830, 2011.

[9] Geoffrey Grimmett and David Stirzaker. *Probability and random processes.* Oxford university press, 2020.

### Supplementary tables

Table 1: Assignment of SNV to nodes of the tree for HGSOC. The first column lists the SNVs and the second column lists the node names.

Table 2: Full list of down regulated pathways in EF compared to ABCD clone (FDR $< 0.01$). The pathways related to the immune system are found to be down regulated. The first column is the name of the gene ontology. The second column is the p-value. The third column is the false discovery rate.

Table 3: Inferred assignment of SNV to nodes of the tree for HER2+. The first column lists the SNV IDs and the second column lists the node names.

Table 4: The hard assignment of cell to nodes of the MAP tree for HER2+ data analysis using PhylEx. The first column is the cell ID and the second column is the node name.

Table 5: Inferred clone fraction for each SNV from the MAP tree for HER2+ data analysis. The first column is the SNV ID and the second column is the clone fraction of the SNV.

150 # Supplementary figures

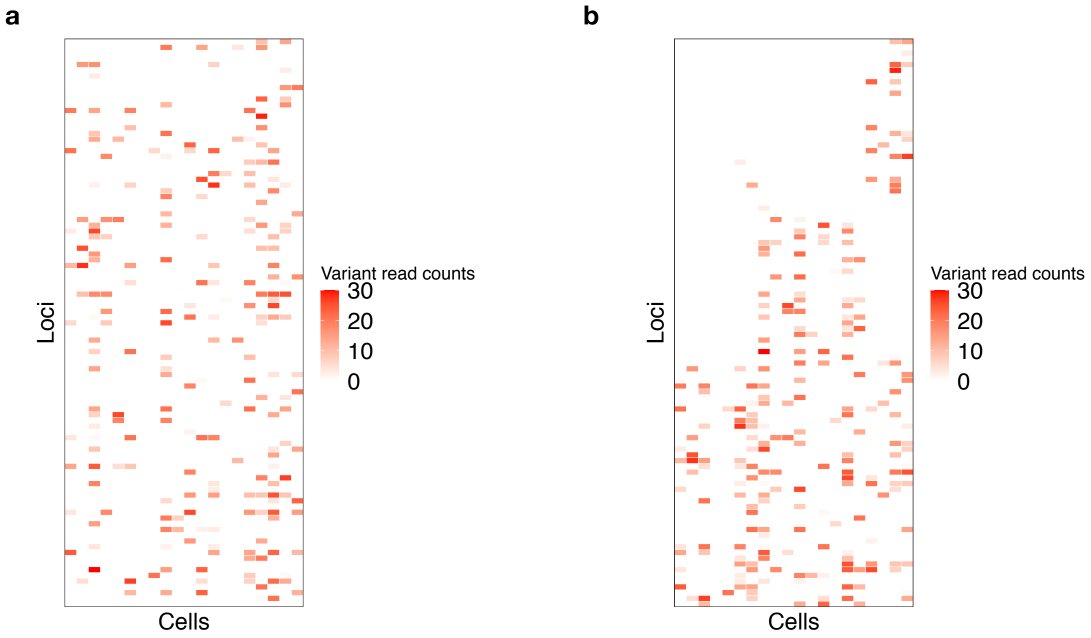


Supplementary Figure 1: Simulated data analysis on a cherry tree. a. Raw gene expression data. b. Gene expression data after co-clustering by SNVs and cells using PhylEx.

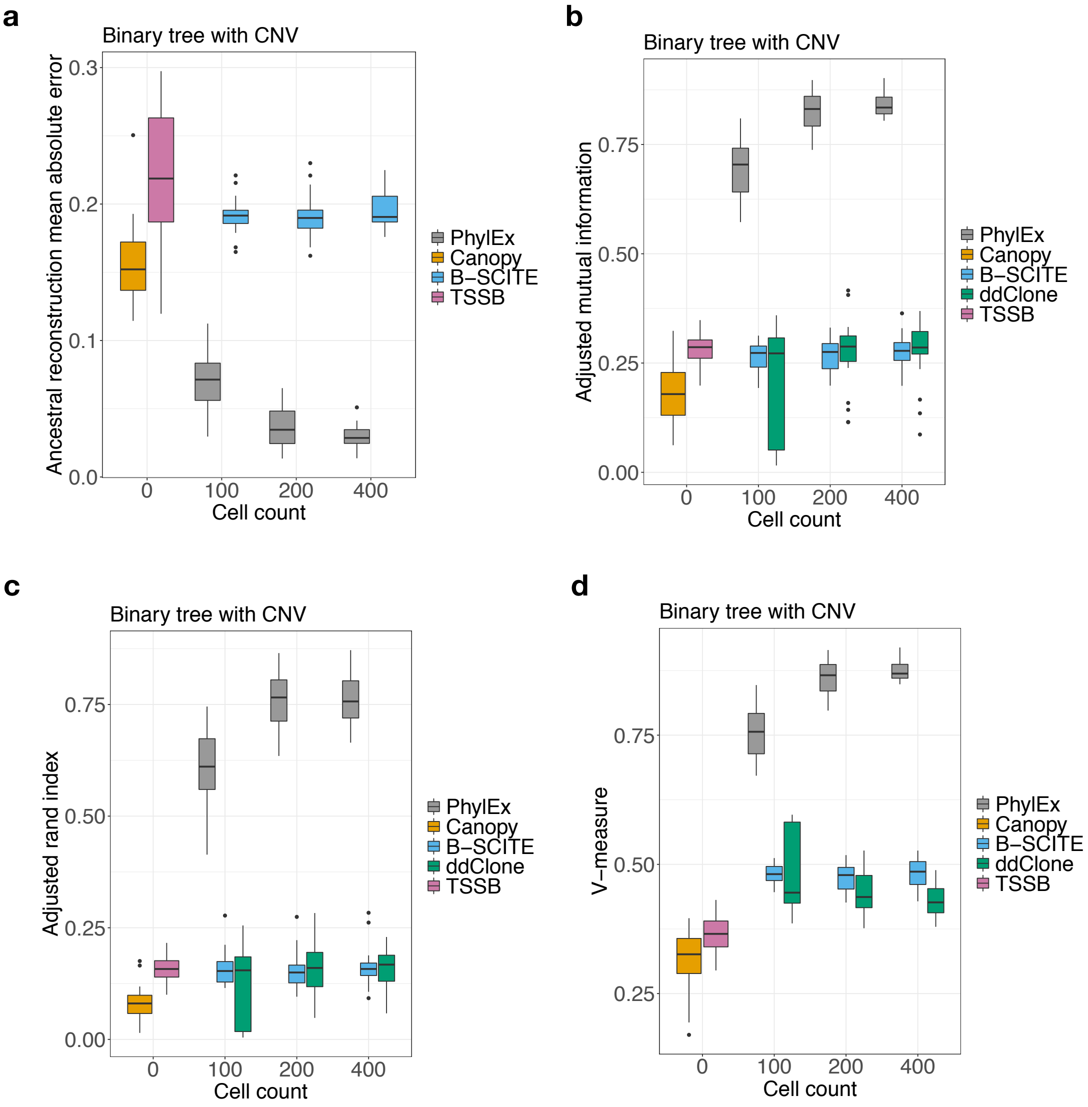


Supplementary Figure 2: Simulated data analysis on binary tree with bulk data generated with copy number evolution on 100 SNVs with $n = 20$ replicates. a. Mean absolute reconstruction error. b. Adjusted mutual information clustering metric. c. Adjusted rand index clustering metric. d. V-measure clustering metric.

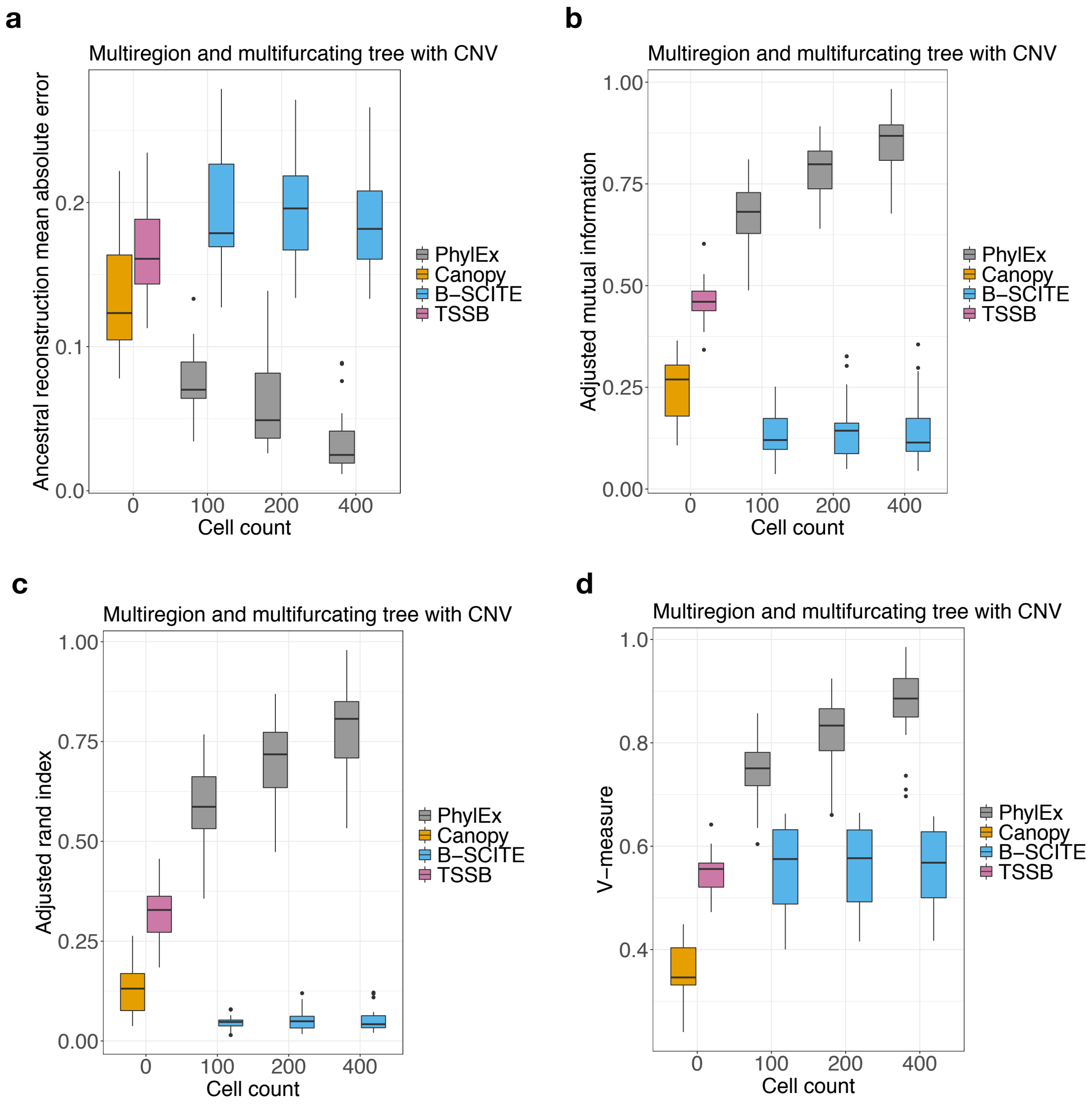


Supplementary Figure 3: Simulated data analysis on multifucating tree with bulk data generated with copy number evolution on 100 SNVs with $n = 20$ replicates. We generated 3 regions to simulate multi-regional bulk data. a. Mean absolute reconstruction error. b. Adjusted mutual information clustering metric. c. Adjusted rand index clustering metric. d. V-measure clustering metric.

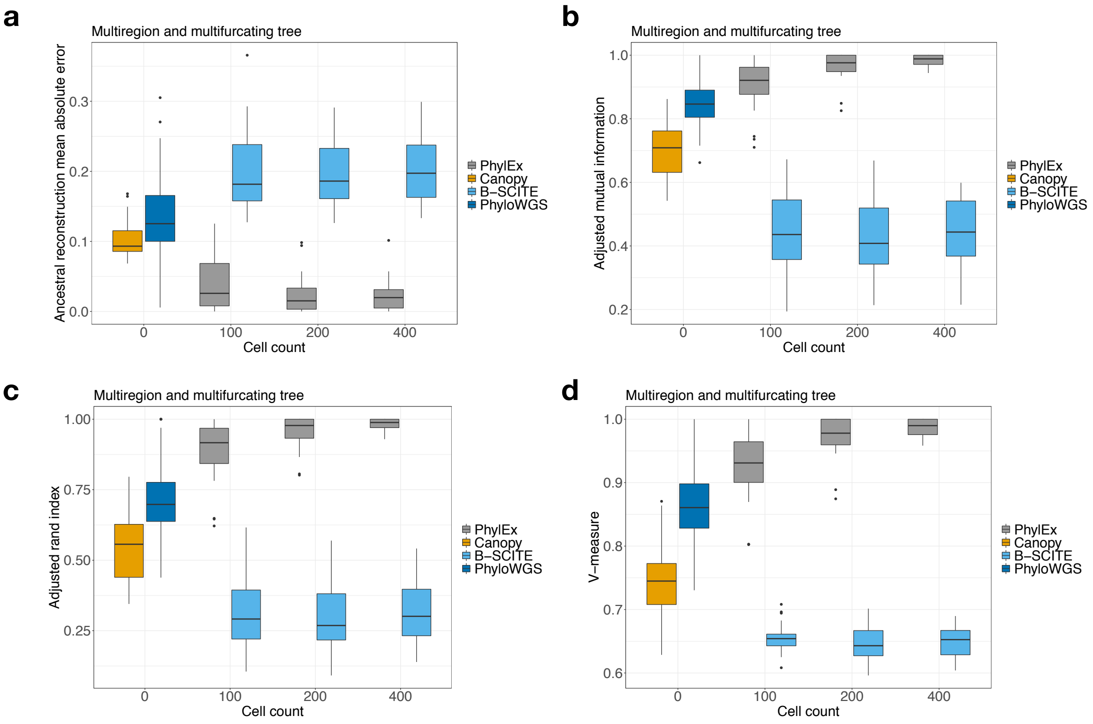


Supplementary Figure 4: Simulated data analysis on multifucating tree with multi-region bulk data generated with copy number evolution on 100 SNVs with $n = 20$ replicates; three regions were generated. The comparison of PhylEx supplied with multi-region and scRNA-seq versus bulk-based methods supplied with multi-region DNA-seq data. a. Mean absolute reconstruction error. b. Adjusted mutual information clustering metric. c. Adjusted rand index clustering metric. d. V-measure clustering metric.

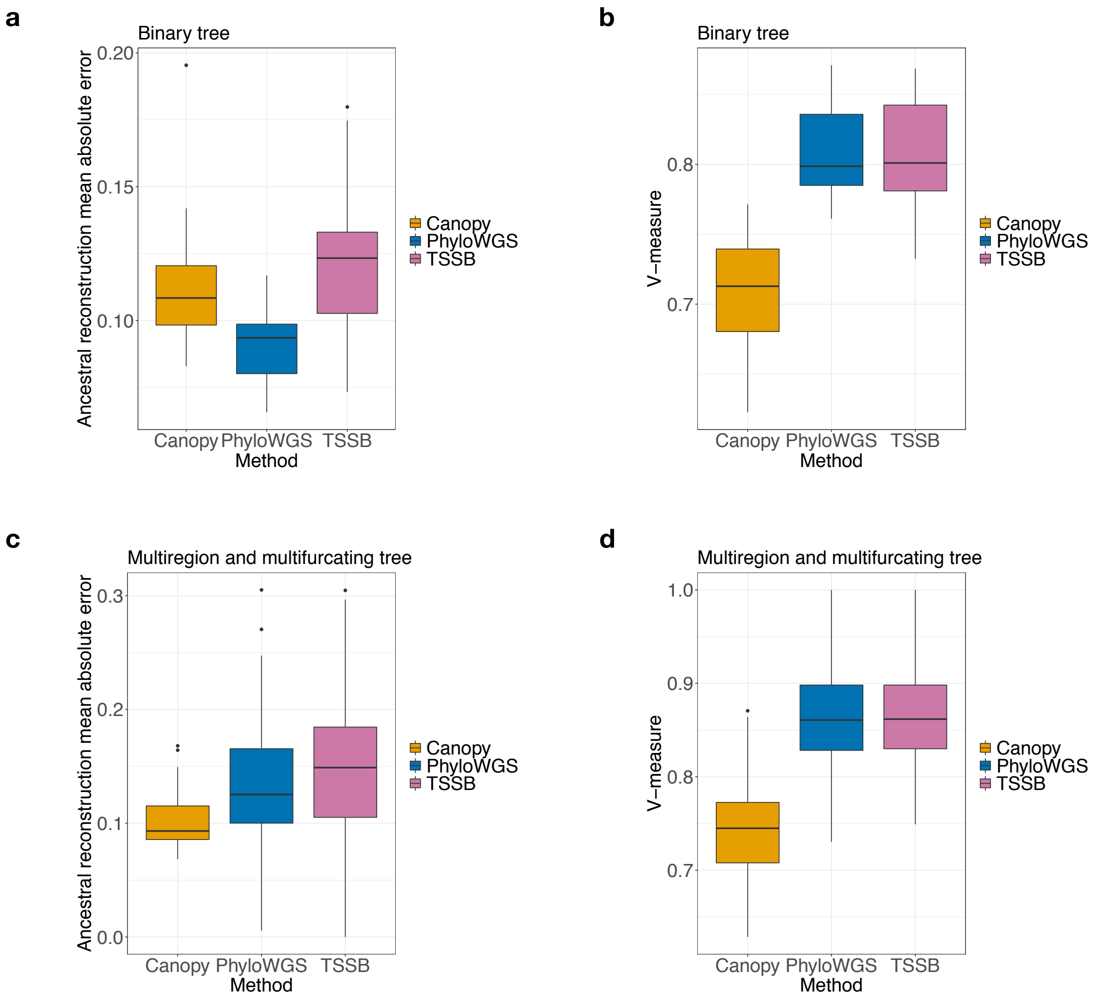


Supplementary Figure 5: The comparison of bulk-based methods on simulated data analysis on binary and multifucating tree with bulk data generated on 100 SNVs with $n = 20$ replicates. Three regions were generated for the multi-region simulation case. The results for the binary tree given single region data is demonstrated using a. ancestral reconstruction error and b. the V-measure clustering metric. The results for multifurcating multi-region data is demonstrated using c. ancestral reconstruction error and d. the V-measure clustering metric. The performances of TSSB and PhyloWGS are on par, allowing us to replace PhyloWGS by TSSB in simulation studies.

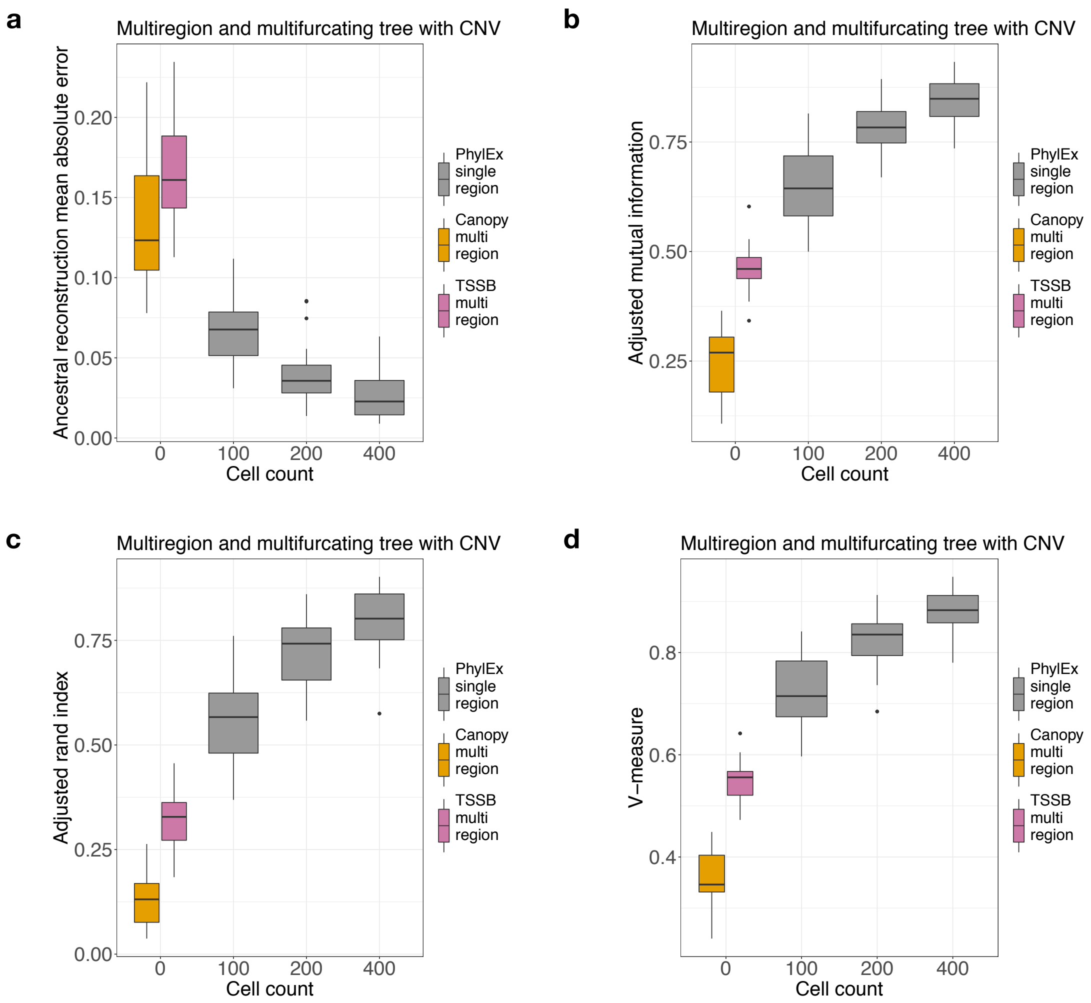


Supplementary Figure 6: Simulated data analysis on multifucating tree with multi-region bulk data generated with copy number evolution on 100 SNVs with $n = 20$ replicates. The comparison of PhylEx using single-region and scRNA-seq versus bulk-based methods supplied with multi-region DNA-seq data. a. Mean absolute reconstruction error. b. Adjusted mutual information clustering metric. c. Adjusted rand index clustering metric. d. V-measure clustering metric. PhylEx using single-region bulk with scRNA-seq data outperforms the bulk methods supplied with multi-region bulk data in all metrics.

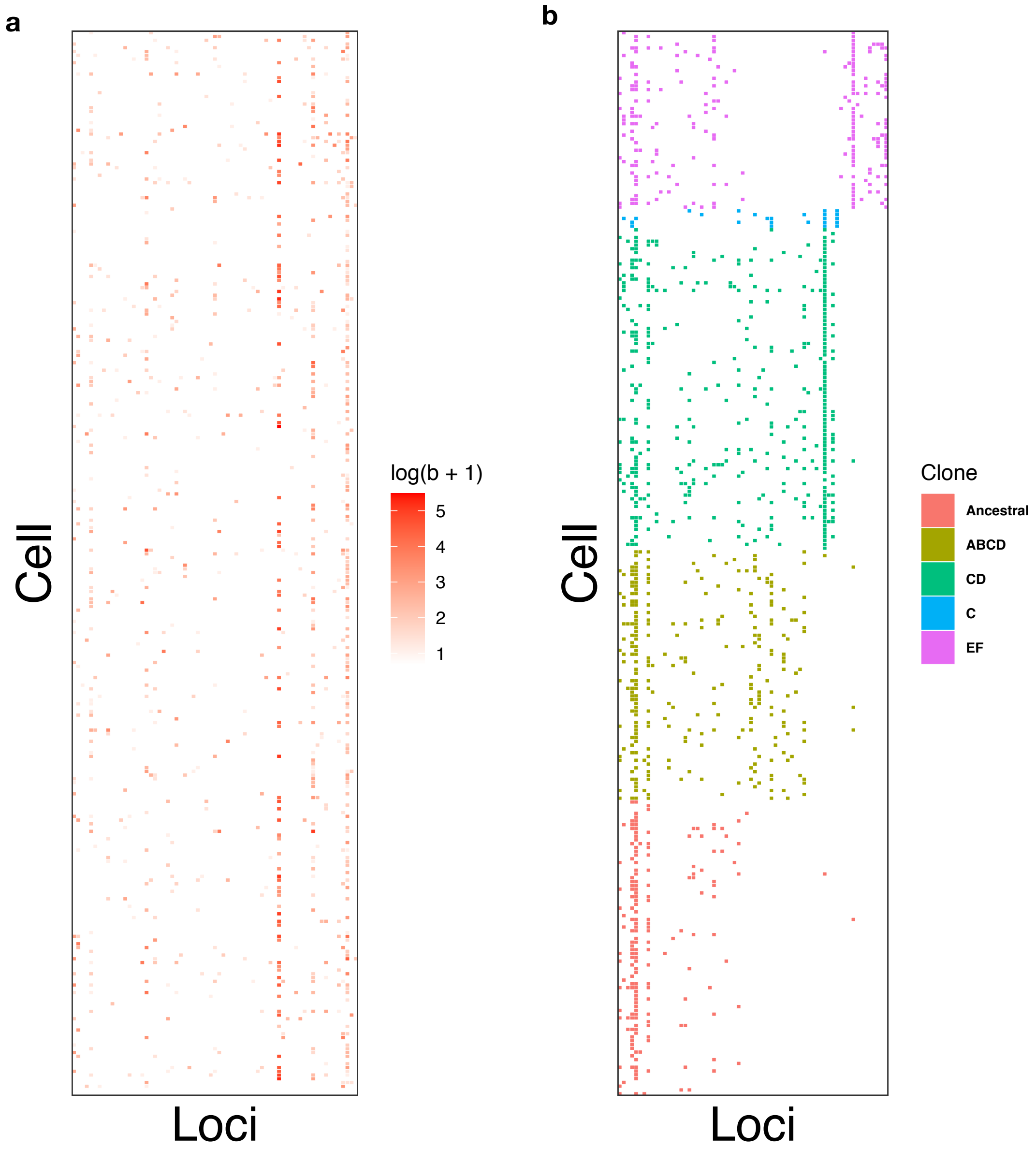


Supplementary Figure 7: a. Raw expression profile in log scale. b. Plot of variant read counts as absence/presence heatmap after co-clustering by cells and SNVs using PhylEx tree.

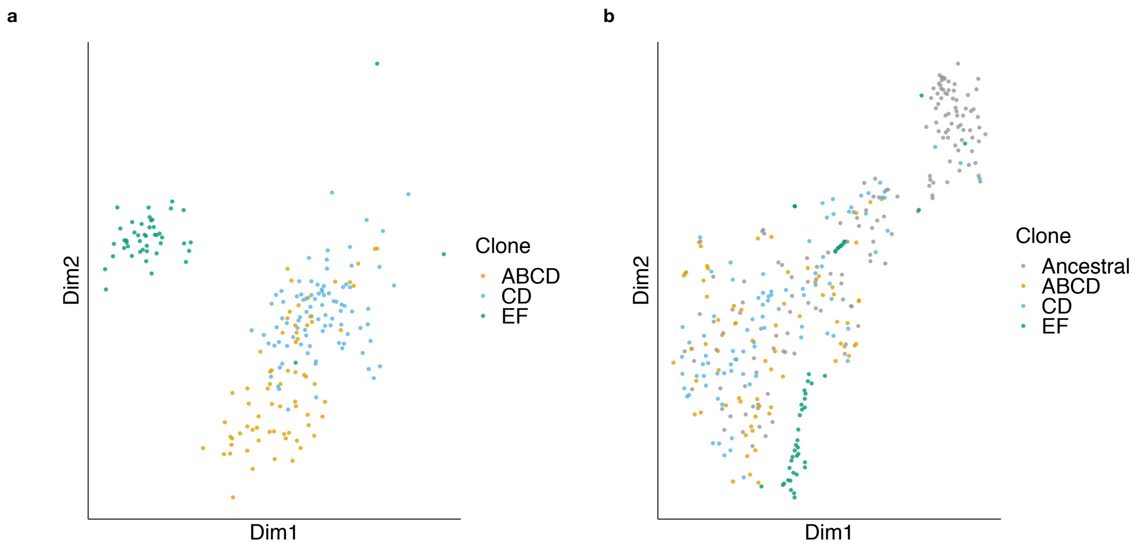


Supplementary Figure 8: a. Plot of gene expressions of cells after performing ZINB-WaVE dimension reduction – plotted without ancestral cells. b. Plot of gene expressions of cells after performing t-SNE dimensions reduction.

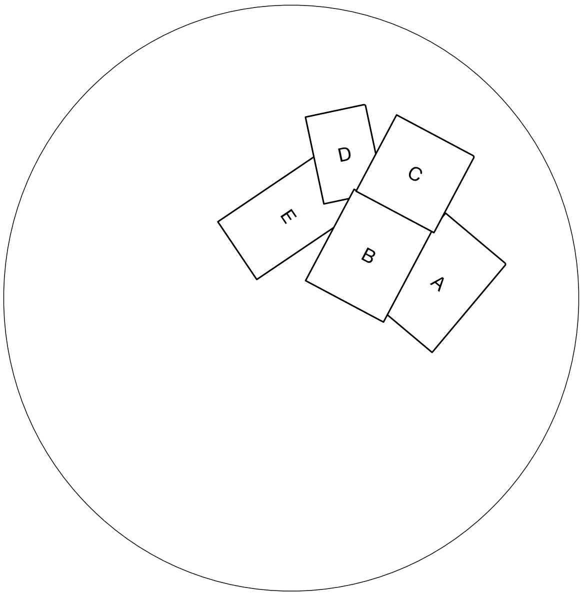


Supplementary Figure 9: Schematic depiction of the locations of multi-regional sampling for HER2+. The proximity of the regions A, B, and C explains the homogeneity in their clone fractions. The region D is remote from region A and the region E does not have any overlap with regions A and C, which explains the differences in the clone fractions in regions D, E to regions A, B, and C.

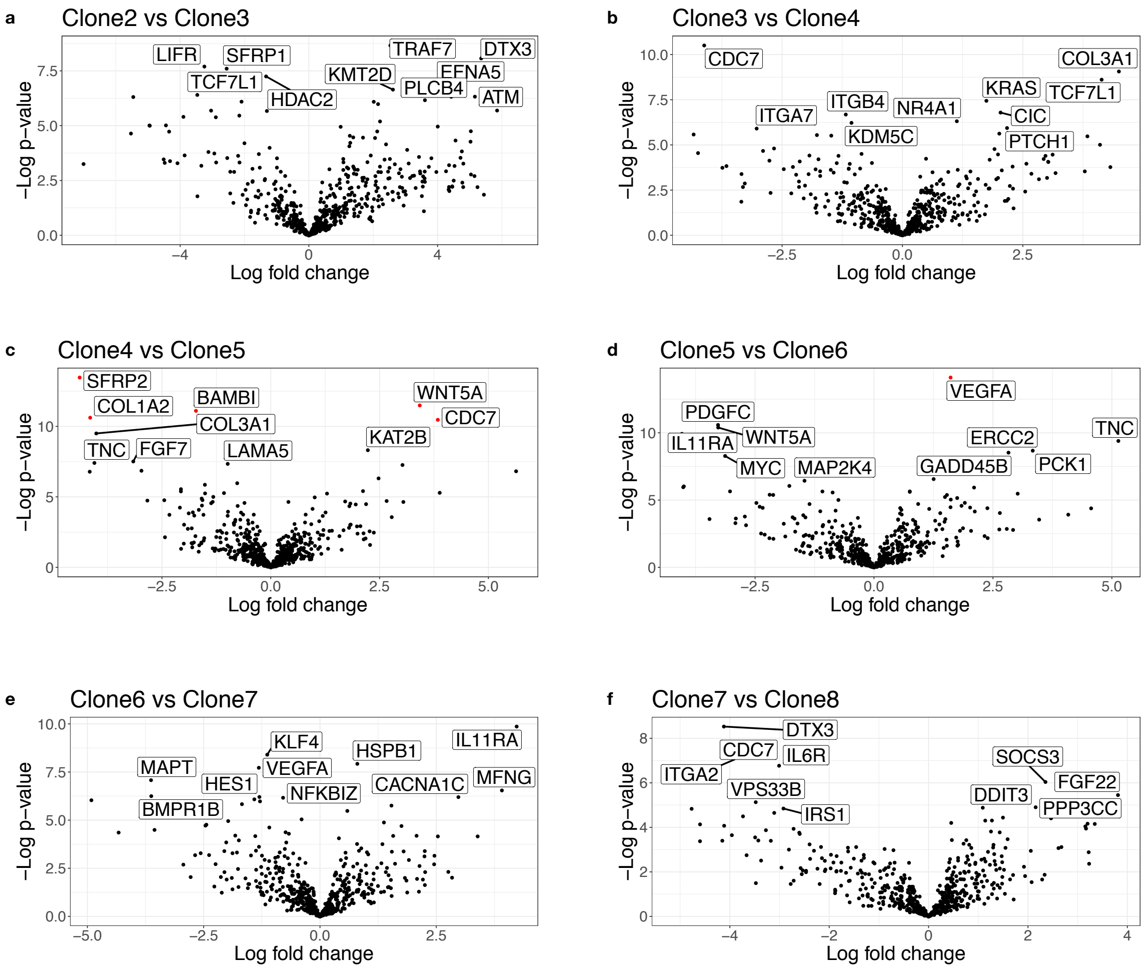


Supplementary Figure 10: a-f. Volcano plots comparing all parent-child clone pairs. Top 10 most significantly differential genes are labelled (FDR $< 0.1$).

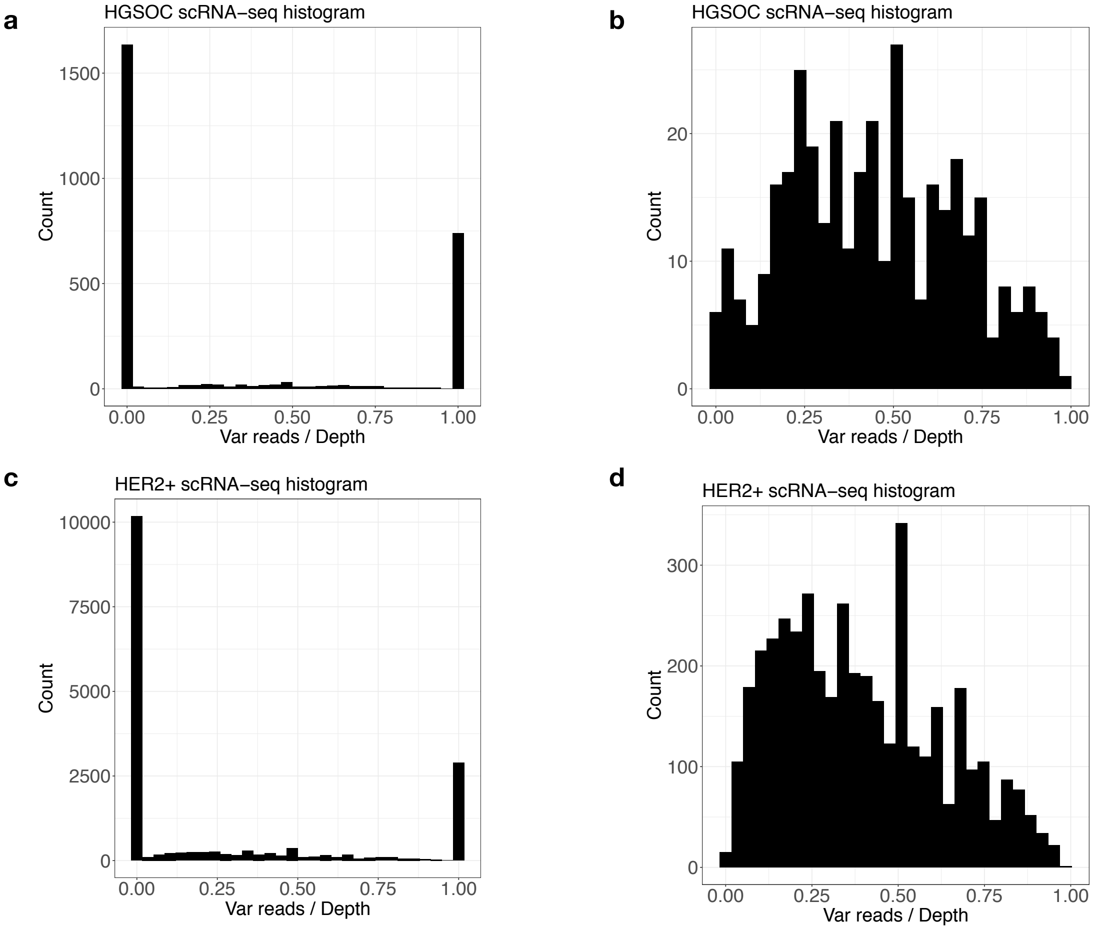


Supplementary Figure 11: Plot of scRNA-seq reads for HGSOC and HER2+ data. a,c. Plot of the ratio of variant reads to depth for scRNA-seq across all loci. b,d. Plot of the ratio of variant reads to depth for scRNA-seq for subset of the data without the extremes at 0 and 1.

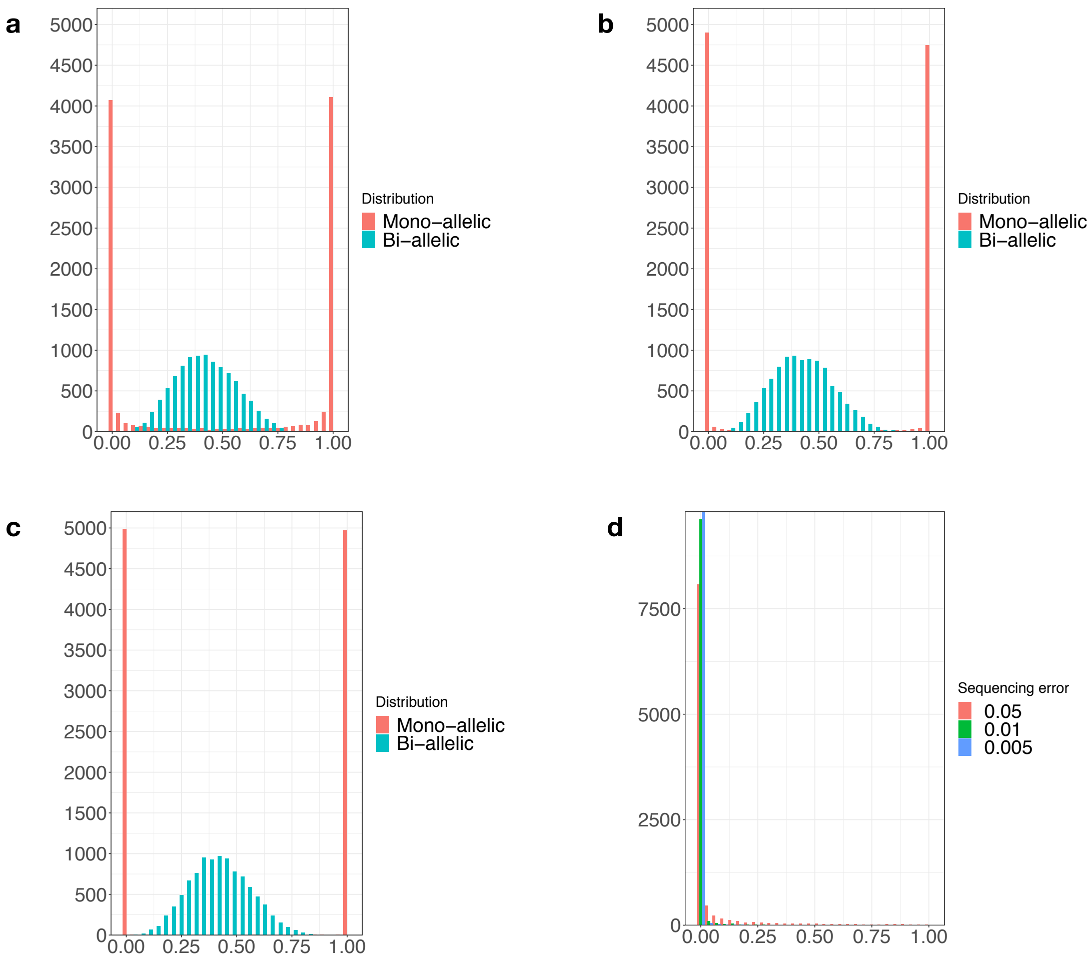


Supplementary Figure 12: a. Beta-Binomial mixture distribution with $\alpha_0 = \beta_0 = 0.05$, b. $\alpha_0 = \beta_0 = 0.01$, c. $\alpha_0 = \beta_0 = 0.005$; the values for the biallelic hyper parameters supplied are $\alpha_n = 5, \beta_n = 7$. As $\alpha_0, \beta_0$ decreases, the mass in the monoallelic distribution concentrates near 0 and 1. d. The ratio of variant reads to depth for the error distribution for $\epsilon \in \{0.05, 0.01, 0.005\}$.

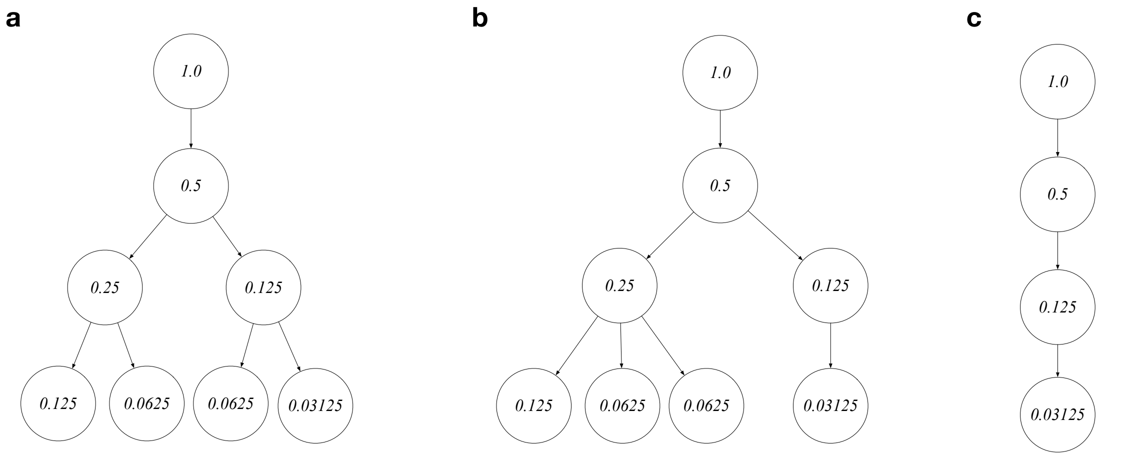


Supplementary Figure 13: a. Binary tree used in the simulation. b. An example of a multifurcating tree used in the simulation. c. A linear tree used in the simulation.
